# Insertion of *YFP* at *P5CS1* and *AFL1* shows the potential, and potential complications, of gene tagging for functional analyses of stress-related proteins

**DOI:** 10.1101/2022.03.08.483394

**Authors:** Toshisangba Longkumer, Louis Grillet, Hao-Yi Chang, Tài Chiến Lường, Chih-Yun Chen, Hadi Putra, Wolfgang Schmidt, Paul E. Verslues

## Abstract

Crispr/CAS9-enabled homologous recombination to insert a tag in frame with an endogenous gene can circumvent difficulties such as context-dependent promoter activity that complicate analysis of gene expression and protein accumulation patterns. However, there have been few reports examining whether such Gene Targeting/Gene Tagging (GT) can alter expression of the target gene. The enzyme encoded by Δ*^1^-pyrroline-5-carboxylate synthetase 1* (*P5CS1*) is key for stress-induced proline synthesis and drought resistance, yet its expression pattern and protein localization have been difficult to assay. We used GT to insert *YFP* in frame with the 5’ or 3’ ends of the endogenous *P5CS1* and *At14a-Like 1* (*AFL1*) coding regions. Insertion at the 3’ end of either gene generated homozygous lines with expression of the *gene-YFP* fusion indistinguishable from the wild type allele. However, for *P5CS1* this occurred only after selfing and advancement to the T_5_ generation allowed initial homozygous lethality of the insertion to be overcome. Once this was done, the GT-generated P5CS1-YFP plants revealed new information about P5CS1 localization and tissue-specific expression. In contrast, insertion of *YFP* at the 5’ end of either gene blocked expression. The results demonstrate that GT can be useful for functional analyses of genes that are problematic to properly express by other means but also show that, in some cases, GT can disrupt expression of the target gene.

**Summary statement:** Gene tagging of *Arabidopsis thaliana P5CS1* and *AFL1* shows the potential of GT for functional analysis of stress-related genes, but also provides examples of how GT can dramatically disrupt expression of the target gene.

---

Proline metabolism plays several, seemingly contrasting roles in stress resistance. Proline accumulation is important for osmotic adjustment, redox regulation and protection of cellular structure (Sharma et al., 2011; Alvarez et al., 2022). However, the combination of high proline synthesis and high rates of proline catabolism is associated with cell death during pathogen infection (Fabro et al., 2004; Senthil-Kumar et al., 2012). Thus, understanding how proline metabolism is regulated is a key topic for many branches of plant stress biology (Verslues et al., 2023). The rate-limiting enzyme of stress-induced proline synthesis is Δ^1^-pyrroline-5-carboxylate synthetase (P5CS), which converts glutamate to the intermediate glutamic semialdehyde, which spontaneously cyclizes to Δ^1^-pyrroline-5-carboxylate (P5C). *P5CS1* is considered to be a canonical stress-induced gene whose expression is used as a marker of stress response in studies too numerous to list here. Despite the hundreds of reports documenting stress-induced proline accumulation and *P5CS1* expression, many questions remain about P5CS1 at the protein level. Particularly, there is a lack of consensus about P5CS1 subcellular localization and how (or whether) it changes in response to abiotic stress (Szekely et al., 2007; Funck et al., 2020). There is also surprisingly limited information on tissue-or cell type-specific accumulation of P5CS1.

For research with Arabidopsis and other model plants, the usual way to answer such questions about subcellular localization and tissue-specific expression is to construct transgenic plants that express the protein of interest, under control of its native promoter and with a fluorescent tag attached, and then use such construct to complement a knock-out mutant. While this approach works well for most genes, it can be problematic because some promoters may not work properly when removed from their normal genomic context. There can also be uncertainty about the length of promoter fragment to use, variability among transgenic lines and instability of transgene expression across multiple generations. The *P5CS1* promoter is heavily methylated (Niederhuth et al., 2016) and promoter methylation is involved in determining stress-induced *P5CS1* expression (Feng et al., 2016). It is unclear whether the proper level of *P5CS1* promoter methylation would be reproduced when the *P5CS1* promoter is inserted into a different genomic context by transformation. Thus, *P5CS1* is a likely example of a promoter whose activity is dependent upon its genomic context. This raises doubt about the use of traditional transgenic reporters to assay *P5CS1* expression patterns.

Gene Targeting, also known as Gene Tagging (here abbreviated as “GT” to encompass both terms), can offer a way around these problems. GT uses homologous recombination to insert or replace sequence at a specific location in the genome (Cermak 2021; Collonnier et al., 2017; Endo et al., 2016; Huang and Puchta, 2021; Huang and Puchta, 2019; Mao et al., 2019; Chen et al., 2022). For example, GT can be used to insert sequence encoding a fluorescent protein or epitope tag in frame with the endogenous protein coding sequence so that the fusion protein can be expressed from its native promoter and in its native genomic context. This may be expected to provide an accurate and stable view of the expression patterns and localization of the protein under study. The first hurdle to feasible GT is the low frequency of homologous recombination in plants and the need the generate a DNA double strand break in the desired location to induce homologous recombination-mediated repair. The limitation on generating site-specific double strand breaks can be overcome by use of Crispr/CAS9-mediated gene editing (Mao et al., 2019; Chen et al., 2022). However, most double-strand breaks are repaired by Non-Homologous End Joining (NHEJ; Huang and Puchta, 2021; Shen et al., 2017). Thus, generation of the double strand break must be coordinated with introduction of a donor DNA fragment containing sequence homologous to the region around the Crispr/CAS9-generated break site such that at least some sites are repaired by recombination rather than NHEJ. It is also critical that homologous recombination events that do occur are heritable. Heritable GT can be enhanced by specifically expressing CAS9 in egg cells or embryos (Wang et al., 2015; Wolter et al., 2018). Note that while we focus here on procedures involving double strand breaks and homology-directed repair, recent work has also reported GT methods that utilize NHEJ to insert new sequence (for example Cermak, 2021; Kumar et al., 2023)

Two types of procedures based on homology-directed repair have been shown to produce relatively high frequencies of heritable GT events in Arabidopsis. The first of these is a two-stage procedure where a transgenic line expressing CAS9 (without guide RNA or with a control guide RNA unrelated to the GT site of interest) is produced. This line is then retransformed with a construct containing the sgRNA and a donor fragment for homologous recombination (Endo et al., 2016; Miki et al., 2018). The donor DNA fragment contains a sequence to be inserted flanked by homology arms which flank the site targeted by the sgRNA to enable homologous recombination once a double strand break has been introduced. After the second transformation, pools of T_2_ plants are screened by PCR to detect the inserted DNA fragment. Individual plants from the positive pools are then screened by PCR and positive plants are backcrossed to segregate out the CAS9 and sgRNA/homology donor constructs. Miki et al. (2018) concluded that the relative success of their strategy was due to the CAS9 being pre-expressed and thus ready to direct DNA cleavage at the moment when the sgRNA/homology donor construct entered the egg cell. They also found that the choice of promoter for CAS9 expression also influenced GT frequency, with DD45 being the best of several promoters tested.

The second strategy is a refined single-stage approach where all components are introduced simultaneously by *Agrobacterium* transformation, but with a system to excise the donor DNA fragment from the larger T-DNA construct (Schiml et al., 2014) or with a translational enhancer to induce higher and more rapid CAS9 protein accumulation (Peng et al., 2020). In single stage GT, it was reported that use of the egg cell-specific promoter *EC1.1* could enhance heritable GT (Wolter et al., 2018). Other experiments revealed that co-expression of chromatin remodelers with the GT construct (Shaked et al., 2005), deletion of a DNA polymerase theta involved in NHEJ (van Tol et al., 2022), heterologous expression of recombinases to promote homologous recombination (Barakate et al., 2020), use of viral replicon expression system to generate increased levels of donor DNA and CAS9 (Cermak et al., 2015; Vu et al., 2020), or use of other nucleases such as CAS12a (Huang et al., 2021; Merker et al., 2020) could enhance GT at different test loci assayed in each study. While these studies all reported ways to enhance GT, they differed with respect to the tested loci, whether the GT test involved insertion of a new sequence (such as YFP or RFP) or just replacement of existing sequence to correct or introduce a readily assayed mutation, and the length of homology arms used in the DNA donor constructs.

Despite the new prospects for efficient GT illustrated in these studies, there is surprisingly little data in plants about the feasibility of GT insertion of a fluorescent tag as an alternative approach to assay patterns of protein accumulation and subcellular localization (Cermak, 2021; Chen et al., 2022). Particularly, there is little data of whether the GT procedure itself can affect expression of the targeted gene. Indeed, some GT studies were in fact designed to perturb expression of the target gene (for example Cermak et al., 2015; Kumar et al., 2023). Overall, there are few examples of GT conducted in Arabidopsis (or other model system) at loci where it is feasible to make a detailed assessment of whether GT altered expression of the target locus or altered the amount or function of the tagged protein produced.

We used GT to insert sequence encoding YFP in frame with either the 5’ or 3’ end of the *P5CS1* coding region. *P5CS1* was a good focus for such experiments because altered expression of *P5CS1* leads to readily detectable changes in stress-induced proline accumulation and because we could use P5CS1 specific antisera to compare levels of P5CS1 protein in wild type to P5CS1-YFP fusion protein after GT. In addition, generation of plants with stable expression of fluorescently-tagged P5CS1 would be a unique resource to answer the many outstanding questions about proline metabolism outlined above. We were ultimately successful in isolating a line expressing P5CS1-YFP which was phenotypically indistinguishable from wild type. Analysis of this line revealed constitutively high levels of P5CS1 in guard cells and tissue specific accumulation in root stele and root meristem.

P5CS1 was cytoplasmic, but typically found adjacent to chloroplasts or in cytoplasmic aggregates. However, such observations were only possible in T_5_ and T_6_ plants as in earlier generations GT events at the 3’ end of *P5CS1* were homozygous lethal. In contrast, GT insertion of YFP at 3’ end of At14a-Like 1 (AFL1), another stress-regulated gene, occurred at high frequency and homozygous GT lines were recovered immediately after transformation with the donor construct. *P5CS1* and *AFL1* were similar in that insertion of YPF at the 5’ end of the coding region abolished expression, for reasons that are unclear. These results show that GT can affect expression of the target gene in ways that vary between loci. Overall, our data illustrate both potential benefits and potential complications of using GT to study gene and protein function. Our generation of a stable P5CS1-YFP line will be a resource for study of this important, but poorly understood, component of abiotic stress resistance.

## Materials and Methods

### Plant materials

Wild-type *Arabidopsis thaliana* (Col-0) and other knock-in or mutant plants were grown on soil at 22°C in a growth room with 16 h light and 8 h dark cycle. Plate-based experiments for control and low water potential (ψ_w_) stress were carried out as described previously (Verslues, 2010). Sterilized seeds were sown on half-strength Murashige and Skoog media supplemented with 2 mM MES [2-(N-morpholino) ethanesulfonic acid] buffer (pH 5.7) and 1.75 % agar without any sugar. After stratification for 3 days at 4°C, plates were placed vertically in a growth chamber maintained at 22°C under continuous light at 90–110 μmol photons m^−2^ s^−1^. Seven-day-old seedlings from control plates were transferred either to freshly prepared control agar plates or agar plates infused with polyethylene glycol (PEG)-8000 to impose moderate severity (−0.7 MPa) low water potential stress. Control and stress samples for gene expression, proline or immunoblot assays were collected at 4 days after transfer.

### Generation of the *EC1.1_pro_:hypaCas9* transformation vector

Plant codon-optimized *HypaCAS9* expressed under the *EC1.1* promoter was generated by mutagenesis of the *EC1_pro_:CAS9* sequence from the pHEE401E plasmid (Wang et al., 2015; Addgene Plasmid #71287). Two fragments were amplified by PCR with oligonucleotide primers harboring the *hypaCas9* mutations described in Chen et al. (2017). This set of PCR products overlapped at the mutation site and also overlapped with the backbone of a Gateway entry clone derived from pDONR221 including the attL1 and attL2 recombination sequences. The two PCR fragments and the vector were ligated through a Gibson Assembly reaction. The *EC1.1_pro_:hypaCas9* fragment was subsequently combined with the pOLE1-OLE1-tagRFP entry clone (Shimada et al., 2010) and then cloned into pH7m34GW destination vector (with Hygromycin resistance for plant selection) by gateway LR reaction. All oligonucleotide sequences used for construct generation are given in Supplemental Table SI (overlapping sequences are highlighted in bold and mutations are represented by upper case letters).

### Generation of the sgRNA and donor DNA transformation vectors

The 5’ and 3’ homology arms for targeting GT to *AFL1* and *P5CS1* were amplified from Arabidopsis Col-0 genomic DNA and YFP (mCitrine) was amplified from pEN2-mCitrine. The mCitrine CDS was initially codon-optimized for Arabidopsis using the online tool from Integrated DNA Technologies (https://sg.idtdna.com/pages/tools/codon-optimization-tool), and synthesized and cloned by AllBio company (Taiwan). Donor plasmid constructs were generated by cloning the PCR amplicon into Gateway entry pDONR vectors; the 5’ arm DNA fragment in P4P1r, YFP (mCitrine) in P1P5r, the 3’ arm DNA fragment in P5P2, and the U626p-tRNA-gRNA in P2rP3. These entry clones were then assembled by multi-site Gateway LR recombination into the destination binary vector pB7m34GW (with Basta resistance for plant selection). For cloning sgRNA, the target-specific crRNA sequence (20 bp) was picked manually by looking at the PAM site located close to the start codon (for GT at the 5’ end of the gene) or close to the stop codon (for GT at the 3’ end of the gene). A pair of sgRNA, having the same target sequence, were cloned for each gene. A pair of reverse complementary primers containing 20 bp of target sequence and an additional flanking 5 bp containing BsaI sites were annealed and digested with BsaI. The digested DNA was mixed with tracRNA and pre-tRNA adapters that had been excised from p1-tRNA-gRNAopti-CmR at the BsaI site, and ligated to pEN-R2-U626p-tRNA-gRNAopti1xAmp-L3 which had also been digested by BsaI (Xie et al., 2015). The sgRNA target sequence and primers used for the construct generation are given in Supplementary Table SI.

### Transgenic line generation and genotyping

To generate the *hypaCas9*-expressing starter line for GT, the *EC1.1_pro_:hypaCas9* construct was transformed into *Arabidopsis thaliana* (Col-0) by the Agrobacterium-mediated floral dipping method. T_1_ seeds were selected on Hygromycin plates and the resistant plants transplanted to soil for generation of T_2_ seeds. T_2_ lines were initially selected on Hygromycin plates. For lines exhibiting 3:1 (resistant:susceptible) segregation ratios consistent with a single locus T-DNA insertion, seedlings were transplanted to soil to collect floral buds for qPCR assay of *hypaCas9* expression. To assay *hypaCas9* expression in subsequent generations, homozygous T_3_ lines from lines T_2_-1 and T_2_-7 were identified by hygromycin selection, and plants from the selected T_3_ lines as well as subsequent T_4_ and T_5_ plants were grown on soil and floral buds collected for quantitative reverse transcriptase-PCR (qPCR) assay.

For the second stage transformation, the vector containing the sgRNA and donor DNA construct was transformed into hygromycin-resistant T_2_ plants, or into homozygous T_3_ plants of the *hypaCas9*-expressing starter line (line #5 and #7). Transformants were selected by spraying Basta herbicide solution on soil-grown plants. Seed of the T_2_ generation (after retransformation) was harvested from Basta-resistant T_1_ plants. Pools of around 30 T_2_ seeds from an individual T_1_ plant were sown on plates without any antibiotics for bulk PCR screening of seedlings. For screening of individual plants from selected lines, T_2_ plants were grown on soil and leaves of individual plants harvested for genomic DNA extraction. Subsequently, T_3_ plants homozygous for the GT-generated YFP insertion were backcrossed to Col-0, and lines homozygous for the YFP insertion but lacking the *EC1.1_pro_:hypaCas9* T-DNA insertion identified by PCR screening. For genotyping of plants after the second transformation, genomic DNA was extracted using the DNeasy Plant Mini Kit (Qiagen). PCR was performed using standard conditions. Primer sequences are given in Supplemental Table SI. T_6_ backcrossed seedlings were used for gene expression and immunoblot experiments except where otherwise noted in figure or figure legend.

### Quantitative RT-PCR analysis of gene expression

RNA of Arabidopsis seedlings or floral buds was extracted the RNeasy Plant Mini Kit (QAIGEN) according to the manufacture’s protocol. RNA amount was quantified using a Nanodrop spectrometer and reverse transcribed using SuperScript III (Invitrogen) and Oligo dT primer. RT-qPCR reactions contained 50 nM cDNA, 200 nM primers and KAPA SYBR FAST master mix (KAPA Biosystems). RT-qPCR reactions and were performed on a 7500 Fast Real-Time PCR System (Thermo Fisher Scientific). *ELF1a* was used as a reference gene to normalize transcript abundances. Primer sequences used for RT-qPCR are shown in Supplemental Table SI.

### Immunoblotting

Protein samples for western blot were prepared by grinding the control and stress-treated seedlings or leaf tissue in liquid nitrogen and suspending the homogenized samples in 200-400 µl of extraction buffer (150 mM NaCl, 50 mM Tris HCl pH 7.5, 10% glycerol, 10 mM EDTA, 10 mM DTT, 1mM NaF, 1% IGEPAL, 2X protease inhibitor, and 2X Roche phosphatase inhibitor) according to the sample quantity (approximately 200 µl of extraction buffer per 100 mg of ground plant material). Protein was quantified using the Pierce™ 660nm Protein Assay Reagent kit (Thermo Scientific). AFL1 and P5CS1 antisera were generated in our laboratory and have been previously characterized (Bhaskara et al., 2015; Kumar et al., 2015), while anti-HSC70 and anti-GFP were obtained from Enzo and Roche, respectively. Immunoblots were performed as previously described (Kesari et al., 2012; Kumar et al., 2015; Wong et al., 2019). In brief, for SDS-PAGE, 10 μg protein was loaded per lane, resolved on 10% polyacrylamide gel, and transferred onto PVDF membrane (4°C overnight, at 30V). The membrane was incubated in 5% milk in TBS-T for at least 1 h at room temperature with shaking. The following dilutions were used for primary and secondary antibodies: anti-Hsc70 (1:5000) /secondary HRP-mouse (1:15000); anti-AFL1 or anti-P5CS1 (1:5000) /secondary HRP-rabbit (1:15000); and anti-GFP (1:5000) /secondary HRP-mouse (1:15000). For anti-GFP, blots were probed with primary antibody overnight at 4°C and for 4 hours at 4°C for the secondary antibody. All other blots were probed for 1 h at room temperature for both primary and secondary antibodies. The signal was detected using the ECL detection kit (Thermo Scientific). Band intensities were quantified using the Fiji (ImageJ) function plot lanes.

### Proline assay

Proline measurement was done using a ninhydrin assay (Bates et al., 1973) adapted to 96-well plate format (Verslues, 2010).

### Confocal imaging

Control and stress-treated seedlings were stained with propidium iodide (PI) to visualize cell walls and then mounted in water or low water potential solution (–0.7 MPa mannitol solution). P5CS1-YFP or AFL1-YFP (excitation at 514 nm and emission at 542 nm for YFP) and PI staining were observed using a Leica Stellaris 8 microscope (PI stained samples were visualized using excitation at 543 nm and emission at 588 nm). For observations of leaf or hypocotyl cells, the Tau-gating function was used to eliminate chlorophyll fluorescence when imaging YFP. Lower magnification images of 3 or 4-day-old seedlings were collected using a fluorescence imager (Zeiss Axio Imager Z1). The tile scan tool was employed to merge several individual images to generate whole-seedling images.

### Statistical Analysis

Data were analyzed by one-factor ANOVA with P-value correction using GraphPad Prism 9 or 10 software. Chi-Square analysis was performed using the GraphPad QuickCalcs online tool.

## Results

### Generation of a starter transgenic line with high level of *hypaCAS9* expression under control of the *EC1.1* promoter

We first transformed wild-type (Col-0) *Arabidopsis thaliana* with a construct for expression of *HypaCAS9* under control of the *EC1.1* promoter. To select a line where CAS9 was efficiently expressed, individual T_2_ lines exhibiting a 3:1 segregation ratio were screened for *hypaCAS9* expression by QPCR. Lines with relatively high expression (line #7 and line #5; Fig. S1A) were used for second stage transformation as described below. We subsequently assayed *hypaCAS9* expression in Line #1 and Line #7 in the T_3_, T_4_ and T_5_ generations to see if CAS9 expression was stable for continued use of these starter lines in subsequent generations. However, in these later generations *hypaCAS9* expression of Line #7 was no longer higher than that of line 1 (Fig. S1B). This may have contributed to lower GT efficiency we observed when using line #7 T_3_ instead of T_2_ as the starter plants for second stage transformation (see below).

### In-frame insertion of *YFP* at the 3’ end of *P5CS1*

For GT at the 3’ end of *P5CS1*, a construct was prepared containing a donor DNA fragment for homologous recombination where YFP was flanked by approximately 1.1 kb homology arms to target recombination to produce in frame fusion of YFP to the P5CS1 C-terminus (Fig. 1A). This length of homology arm was similar to that used in previous studies (Miki et al., 2018; Shaked et al., 2005; van Tol et al., 2022; Wolter et al., 2018). The construct also encoded an sgRNA to direct DNA cleavage next to the *P5CS1* stop codon (Fig. 1A). This cleavage site was not present in the donor fragment because the *P5CS1-YFP* fusion disrupted the sgRNA binding site. Thus, the sgRNA could not direct cleavage of the donor fragment or the endogenous YFP-fused gene after homologous recombination. This construct was then transformed into T_2_ *EC1.1_pro_:hypaCAS9* plants described above (line #5) or into T_3_ of line #7. Retransformed transgenic plants were selected by Basta. For the line #5 transformation, an initial screening of 48 T_2_ pooled DNA extracts (each pool contained seed from one T_1_ Basta-resistant plant, approximately 30 seedlings collected for each pool) identified four positive T_2_ DNA pools (Fig. 1B; Table 1). Screening of individual seedlings from two of these pools identified GT insertion of the donor fragment, as indicated by amplification of PCR products of the expected size using primers specific for *YFP* and for *P5CS1* sequence just outside of the homology arms (primers 2 and 3, Fig. 1A). The frequency of GT insertion we obtained was consistent with that observed in a previous two stage GT procedure (Miki et al., 2018). A similar screening was conducted for the line #7 T_3_ transformation. In that case, the second stage transformation was successful, as indicated by amplification of the higher molecular weight band by primer set 1 showing that the T-DNA containing the donor construct had been inserted into the genome (Supplemental Fig. S2). However, none of the lines had *YFP* inserted at the *P5CS1* locus, as indicated by the lack of amplification from primer sets 2 and 3 (Supplemental Fig. S2B). It is likely that this lack of GT activity was due to the lower level of *hypaCAS9* expression in the line #7 T_3_ generation (Supplemental Fig. S1), since line #7 T_2_ transformation yielded high frequency of YFP insertion at the AFL1 locus (see below).

**Figure 1:**
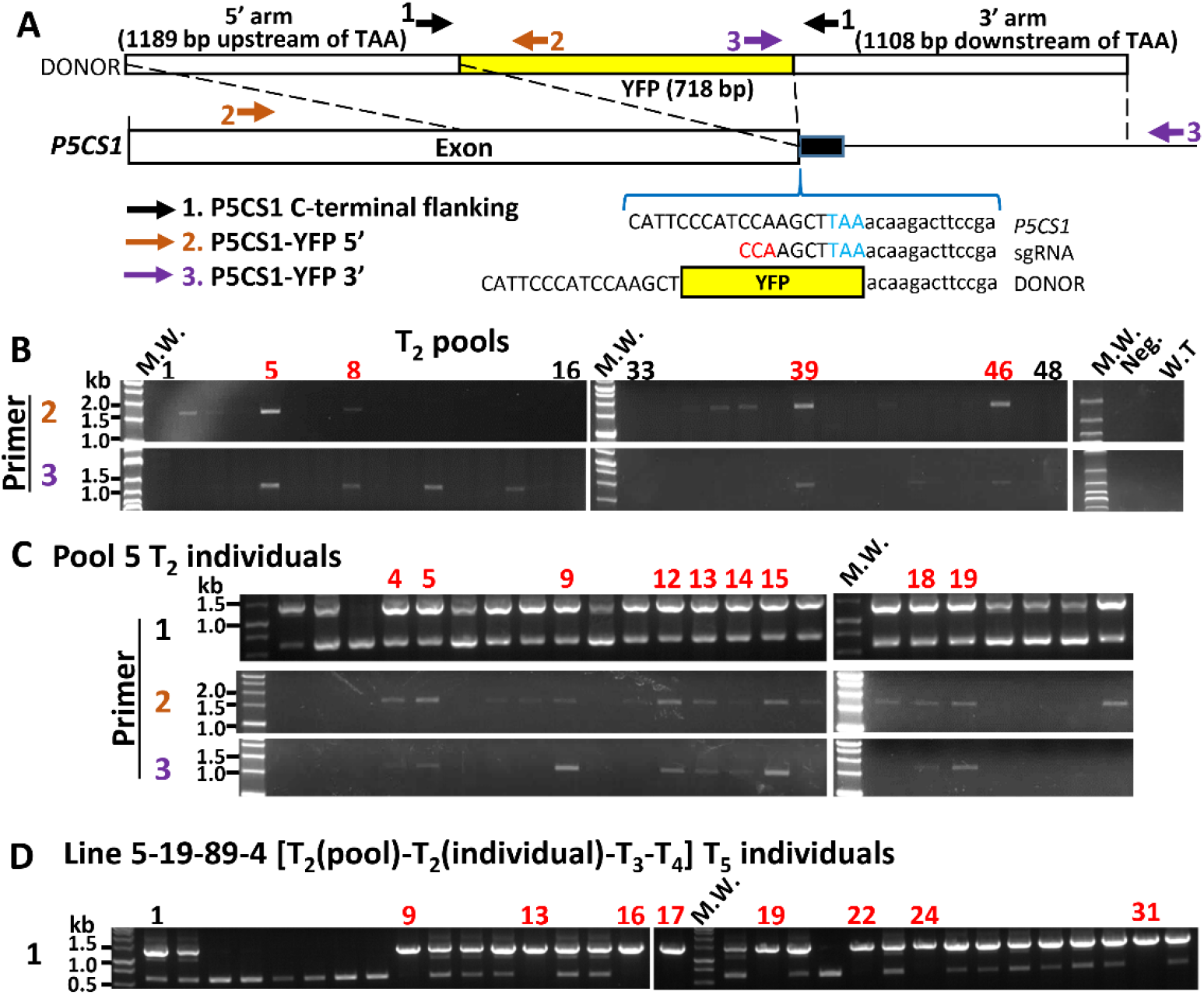
GT insertion of *YFP* at the 3’ end of *P5CS1*. A. Donor Fragment design and genotyping primer positions for knock-in of C-terminal YFP at the *P5CS11* locus. The *YFP* coding sequence was flanked by homology arms matching the indicated regions of *P5CS1*. In the DNA sequences shown, blue indicates the *P5CS1* stop codon and red indicates the PAM site on the sgRNA. Note that replacement of the *P5CS1* endogenous sequence with the knock-in fragment disrupts the sgRNA recognition site thus preventing cutting of the donor fragment or re-cutting of endogenous *P5CS1* after insertion of the donor fragment. Positions of primers used for genotyping are also shown (primer sequences and amplicon sizes can be found in Supplemental Table I). B. Genotyping of pooled T_2_ seedlings from 48 Basta resistant T_1_ plants using primer sets 2 and 3. Red numbers indicate T_2_ seed pools with insertion of the YFP donor construct at the *P5CS1* locus. Note the pools 17-32 did not contain any positives and are not shown. C. Genotyping of individual T_2_ plants from line 5. Red numbers indicate plants containing the *P5CS1-YFP* knock-in based on amplification of the expected size PCR product for primers 2 and 3. Note that no homozygous plants were obtained. D. Genotyping of T_5_ plants from the T_4_ line positive for P5CS1-YFP production (Supplemental Fig. S5). Red numbers indicate plants homozygous for the *P5CS1-YFP* knock-in. Note that the donor construct had already been removed from these plants by genetic segregation in previous generation. Thus, primer set 1 could be used to genotype for the *P5CS1-YFP* knock-in based on presence or absence of the high molecular weight band (from *P5CS1-YFP*) or the lower molecular weight band (wild type *P5CS1*).

**Table 1:** GT genotyping results and efficiency of 5’ and 3’ *YFP* insertion at the *AFL1* and *P5CS1* loci. The number of homozygous, heterozygous and W.T. plants were counted based on the flanking PCR (primer set 1) and the presence bands of expected molecular weight for 5’ and 3’ YFP amplicons (primer sets 2 and 3). Primer positions and PCR results are shown in Figures 1 and 5 as well as Supplemental Figures S2, S9, S12, S13 and S14. Putative single crossovers scored as plants that had amplification of the 5’ or 3’ YFP PCR but not the other side. Asterisks (*) indicate lines where a band of higher than expected molecular weight was observed on 3’ YFP PCR indicating that the construct may have undergone NHEJ incorporating YFP and all (or a portion) of the 3’ homology arm rather than homologous recombination. NA = Not Assayed/Not Applicable

|  | Second Transformation:<br>Bulk Screening of T<sub>2</sub> |  | Second transformation: Screening of Individual T<sub>2</sub> Plants |  |  |  |  |
| --- | --- | --- | --- | --- | --- | --- | --- |
| Parental Cas9 Line # and generation | PCR Positive | GT Efficiency | T<sub>2</sub> line | Null (W.T.) | Heterozygous | Homozygous | Putative Single Crossover |
|  | ***P5CS1* 3' (P5CS1-YFP)** |  |  |  |  |  |  |
| #5 (T<sub>2</sub>) | 4/48 | 8% | 5 | 8 | 8 | 0 | 6 |
|  |  |  | 8 | NA | NA | NA | NA |
|  |  |  | 39 | 8 | 8 | 0 | 7 |
|  |  |  | 46 | NA | NA | NA | NA |
| #7 (T<sub>3</sub>) | 0/40 | 0% | NA | NA | NA | NA | NA |
|  | ***AFL1* 3' (AFL1-YFP)** |  |  |  |  |  |  |
| #7 (T<sub>2</sub>) | 6/30 | 20% | 11 | 7 | 6 | 11 | 7 |
|  |  |  | 14* | 1 | 13 | 10 | 1 |
|  |  |  | 26 | 5 | 9 | 10 | 3 |
|  | ***P5CS1* 5' (YFP-P5CS1)** |  |  |  |  |  |  |
| #7 (T<sub>3</sub>) | 2/30 | 7% | 6 | 4 | 16 | 4 | 2 |
|  |  |  | 13* | 4 | 15 | 5 | 7 |
|  | ***AFL1* 5' (YFP-AFL1)** |  |  |  |  |  |  |
| #7 (T<sub>2</sub>) | 1/29 | 3.5% | 26 | 2 | 17 | 5 | 0 |

We then genotyped individual plants from two selected T_2_ seed pools (Table I, Fig. 1C). In both pools, plants that had undergone homologous recombination were identified based on amplification of the expected size PCR product for primer 2 and primer 3. Sequencing of selected plants confirmed in-frame insertion of YFP (Supplemental Fig. S3). None of these plants were homozygous for the YFP knock-in since primer 1 still amplified the lower molecular weight band from wild type P5CS1. We also observed plants where the correct fragment was amplified at the 5’ side but not at the 3’ side of the YFP insertion. These may be “single crossover” lines where homologous recombination occurred at the 5’ end of the donor construct (based on the primer 2 results) but some other insertion or deletion event, possibly involving NHEJ instead of homologous recombination, occurred at the 3’ end of the donor construct (see *AFL1* section below for further illustration of such putative single cross-over events).

Plants that were heterozygous for the *P5CS1-YFP* knock-in were allowed to self and T_3_ progeny plants genotyped. However, no individuals homozygous for the *P5CS1-YFP* insertion were identified (Supplemental Fig. S4A, C). In addition, siliques of heterozygous plants (containing T_4_ seed) had a high proportion of missing seed, indicative of embryo abortion or defect in seed development (Supplemental Fig. S4B). The segregation ratios of wild type versus heterozygous plants or normal versus aborted seed matched the 2:1 ratio expected if the homozygous *P5CS1-YFP* knock-in was lethal (exemplified by the results of line 5-9 and 5-19; Supplemental Fig. S4A-C). Also, we were unable to detect P5CS1-YFP fusion protein in heterozygous T_3_ plants, suggesting that expression of the knock-in allele had been disrupted (Supplemental Fig. S4D). However, we did identify some T_3_ lines where almost no seed abortion was observed (for example line 5-19-88 and 89, Supplemental Fig. S4E). Individual T_4_ progeny from these two T_3_ plants were then grown and genotyped. We still did not find plants homozygous for the *P5CS1-YFP* insertion, and for some of these plants a high rate of missing seed was still observed (line 5-19-88; Supplemental Fig. S5A). In others, missing seed was not observed (line 5-19-89; Supplemental Fig. S5B). From this line, a heterozygous plant (5-19-89-4) was identified where both P5CS1 and P5CS1-YFP could be detected (Supplemental Fig. S5C). The T_5_ progeny of this plant exhibited the expected 1:2:1 segregation ratio for P5CS1-YFP (Fig. 1D; Chi-square P = 0.91). This T_4_ line was also backcrossed to wild type and homozygous plants recovered in the F_2_ (T_6_) generation were grown to maturity and seed collected for physiological assays.

Although it was not an objective of our study to determine the mechanism underlying this apparent homozygous lethality and later recovery of the *P5CS1-YFP* GT insertion, we do note that there are heavily methylated regions in the *P5CS1* promoter and region downstream of *P5CS1* (Supplemental Fig. S6). Thus, lack of P5CS1-YFP accumulation until the T_4_ generation suggests the hypothesis that GT disrupted the epigenetic state around the *P5CS1-YFP* insertion, which then had to be reset before the GT-modified gene could be expressed. We also do not rule out the possibility that disrupted expression of *AT2G39795*, which is adjacent to the 3’ end of *P5CS1*, could have contributed to the initial homozygous lethality of the *P5CS1-YFP* GT insertion.

### In the T_6_ generation, P5CS1-YFP plants were indistinguishable from wild type in P5CS1 expression and proline accumulation

Progeny of two of the homozygous plants identified in Fig. 1D (5-19-89-4-9 and 5-19-89-4-13) had *P5CS1* gene expression that was indistinguishable from wild type (Fig. 2A). In wild type, *P5CS1* expression was induced nearly 7-fold after transfer of seedlings to −0.7 MPa for 96 h, consistent with previous data from our laboratory. *P5CS1-YFP* plants had wild type levels of proline accumulation in response to low water potential (Fig. 2B), again indicating that *P5CS1-YFP* was expressed normally and that the YFP tag did not interfere with P5CS1 function.

**Figure 2:**
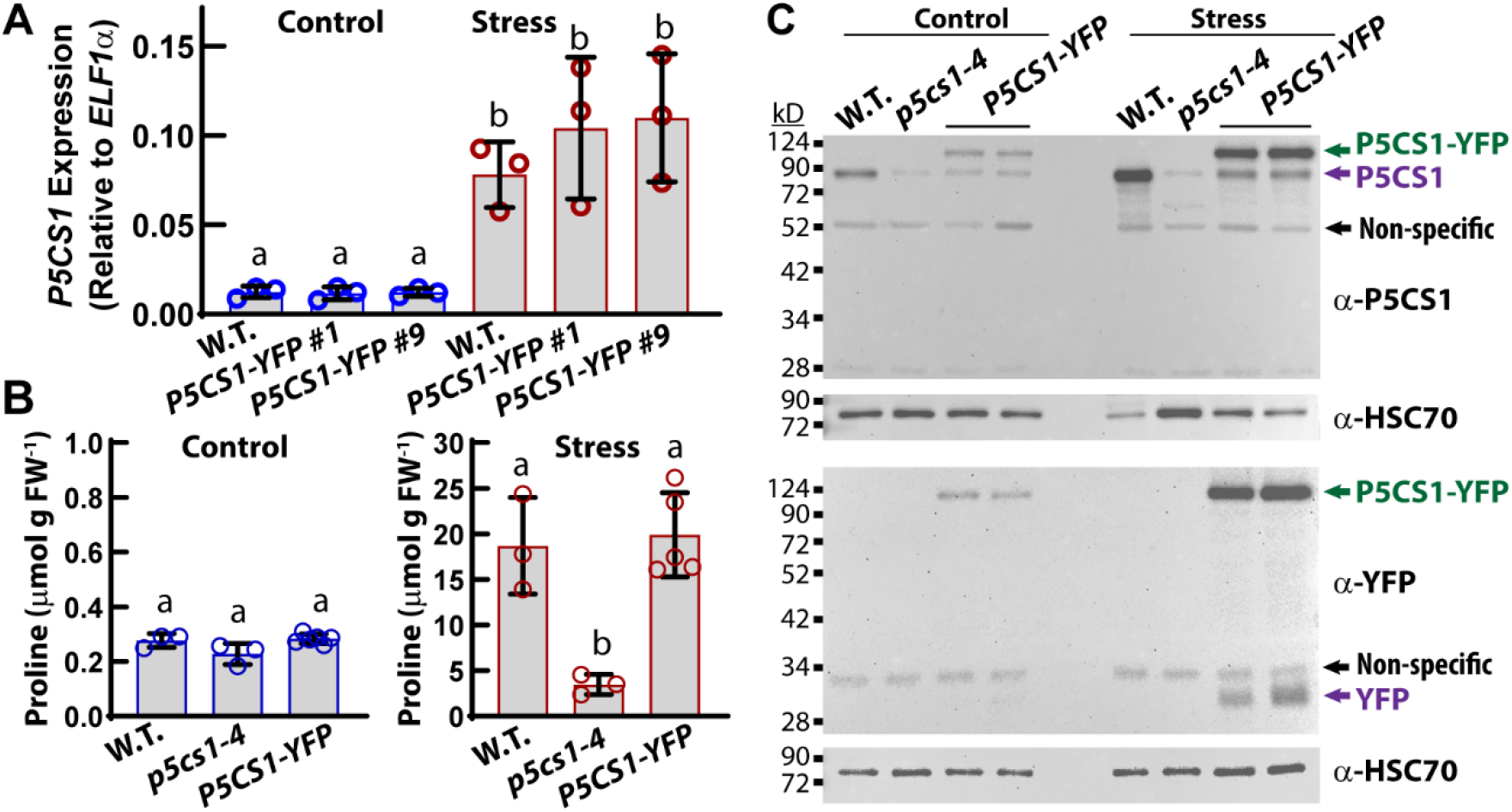
In the T_6_ generation, P5CS1-YFP plants have *P5CS1* gene expression, proline accumulation and P5CS1 protein levels indistinguishable from wild type. A. *P5CS1* gene expression. Seven-day-old seedlings were transferred from control media to either fresh control plates or PEG-infused agar plates (−0.7 MPa) for low water potential treatment. Gene expression is calculated relative to the reference gene *ELF1a* whose expression level is not affected by the stress treatment. Data are means ± S.D. from three independent experiments (n = 3 for wild type). For *P5CS-YFP*, seed from two homozygous plants (5-19-89-4-9 and 5-19-89-4-13 gave indistinguishable results and the combined data are shown (n = 6). Different letters indicate significant difference (ANOVA, correct P B. Proline contents of wild type, *p5cs1-4* and *P5CS1-YFP*. Stress treatments and data format are the same as described for panel A. C. Immunoblot of P5CS1 demonstrates that P5CS1-YFP accumulates to levels similar to wild type in both stress and control treatments. For each lane, 20 μg of protein was loaded and P5CS1 or YFP detected. For *P5CS-YFP*, seedlings from the same two homozygous plants (5-19-89-4-9 and 5-19-89-4-13) used for gene expression and proline determination were used for immunoblot. Detection of HSC70 was performed as a loading control. The expected molecular weight of P5CS1 is 78 kD, although we typically observe that it runs at a slightly higher apparent molecular weight. The molecular weight of YFP plus the linker sequence used to attach it to P5CS1 is 29-30 kD.

Immunoblot using P5CS1 antisera showed that the molecular weight of P5CS1 was shifted up by approximately 30 kD, consistent with the molecular weight of YFP plus the short linker sequence (Fig. 2C). Note that wild type P5CS1 consistently runs higher than its expected molecular weight of 78 kD (Kesari et al., 2012). Comparison to the *p5cs1-4* mutant showed that the band detected at approximately 85 kD is specific to P5CS1, with only very low level of cross reactivity, possibly due to P5CS2 (Kesari et al., 2012). In the P5CS1-YFP samples there still was a low level of protein detected at the apparent molecular weight of P5CS1 alone. Immunoblot detection of YFP showed that P5CS1-YFP plants had a similarly low level of free YFP (Fig. 2C). This indicated that a small portion of P5CS1-YFP was post-translationally cleaved to yield free YFP and P5CS1. Comparison to AFL-YFP generated by the same GT procedure (see below) indicates that this apparent proteolytic removal of the YFP tag is specific to P5CS1 and not related to the GT procedure.

### Guard cells have constitutively high P5CS1 while low **ψ_w_** can induce P5CS1 accumulation in both root and shoot tissues

This successful YFP-tagging of endogenous *P5CS1* revealed new aspects of P5CS1 tissue-specific accumulation. Perhaps most striking was that P5CS1-YFP accumulated in guard cells, but not leaf pavement cells, in both control and low ψ_w_ treatments (Fig 3A, Supplemental Fig. S7, S8). This was consistent with guard cell gene expression profiling which had also found high expression of *P5CS1* (Supplemental Fig. S8A) (Yang et al., 2009). P5CS1-YFP was constitutively present in hypocotyl epidermis and leaf mesophyll cells as well as the root meristem region and the tip of root hairs (Supplemental Fig. S7A). During low ψ_w_ stress, P5CS1 levels were increased in the root tip, mature root, hypocotyl, and leaf (Supplemental Fig. S7B). Higher resolution imaging found that in response to low ψ_w_ stress, P5CS1-YFP increased in all cell layers of the root meristem and elongating cells, while in more mature sections of the root P5CS1-YFP accumulated specifically in the steele (Fig 3B). The accumulation of P5CS1 in cortex cells of root tip was consistent with previous sequencing data (Fig. S8C). During low ψ_w_, P5CS1-YFP did not accumulate in the root quiescent center, center of the shoot meristem, or leaf pavement cells (Fig. 3, Supplemental Fig. S8). These data demonstrated that while *P5CS1-YFP* was widely expressed, especially at low ψ_w_, it also maintained a degree of tissue specificity.

**Figure 3:**
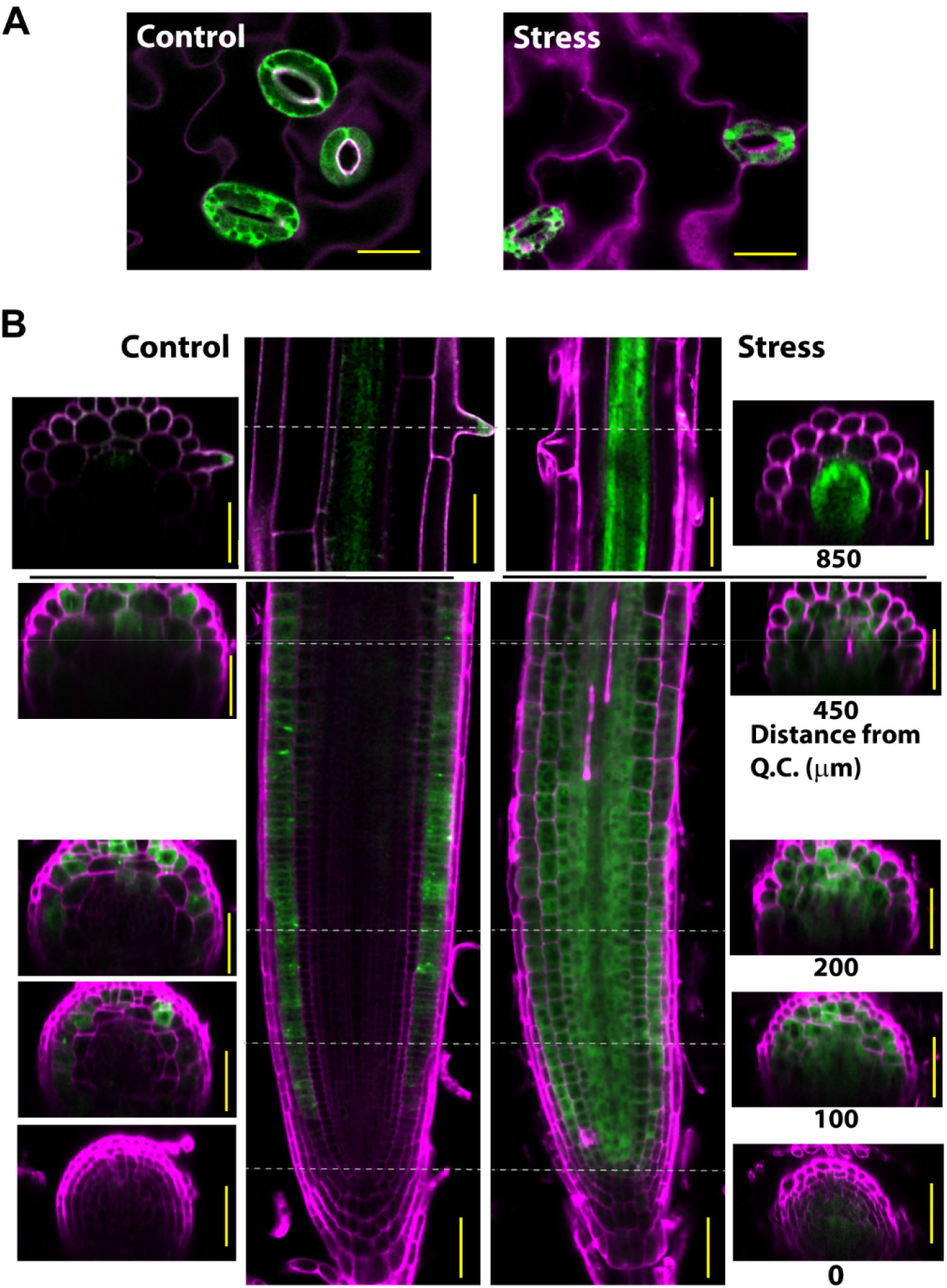
P5CS1-YFP knock-in reveals tissue specific patterns of P5CS1 accumulation. A. P5CS1-YFP accumulates to high levels in guard cells under either control or stress conditions but is not detectable in leaf pavement cells. Scale bars indicate 20 μM. Data are presentative of multiple plants and multiple experiments. B. Pattern of P5CS1-YFP accumulation in the root tip and elongating cells. In the unstressed control P5CS1-YFP is present at relatively low level in epidermal and cortex cells and sometimes forms foci within these cells. Note also the presence of P5CS1 in the tip of growing root hairs. During low ψ_w_ stress (−0.7 MPa) P5CS1-YFP accumulates across all cells in the root meristem (but not quiescent center) and is present at high level in the steele of the root elongation zone and much of the mature root. Numbers under the cross section images on the stress side indicate the distance from the quiescent center. Cross sections from the unstressed root are from the same positions. Scale bars indicate 50 μM. Data are representative of images collected from multiple plants grown in multiple independent experiments.

Higher resolution imaging of leave mesophyll cells and hypocotyl cells, using Tau-gating to eliminate background chlorophyll fluorescence from the YPF channel, found that P5CS1-YFP was cytoplasmic under both control and stress conditions (Fig. 4A and B). That said, P5CS1 was predominantly localized around the outside of the chloroplast and high levels of P5CS1-YFP were often observed between two or more closely spaced chloroplasts (Fig. 4). This contrasted with root cells, where P5CS1-YFP was diffusely localized in the cytoplasm (Fig. 3B). The proximity of P5CS1 to chloroplasts is of interest given that proline synthesis has been hypothesized to help balance chloroplast redox status during low ψ_w_ stress (Shinde et al., 2016; Zheng et al., 2021). We also found that P5CS1-YFP was occasionally observed in aggregates in unstressed root (Fig. 3B). These may be similar to the P5CS1 aggregates described by Szekely et al. (2008); although, the precise nature of such aggregates is unknown. Such aggregates were not observed in plants at low ψ_w_ despite the overall higher level of P5CS1 accumulated during low ψ_w_ stress.

**Figure 4:**
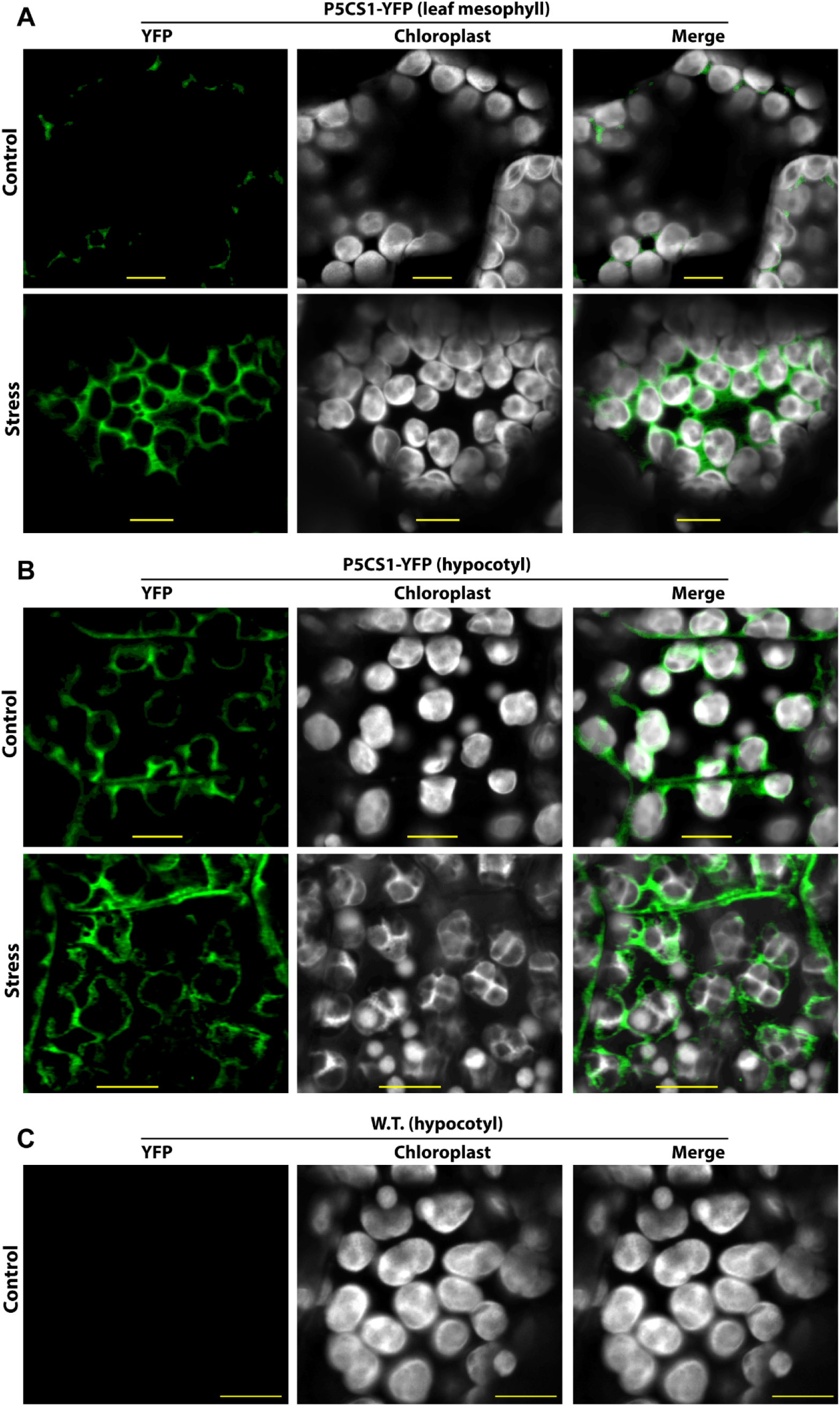
Subcellular localization of P5CS1-YFP in leaf mesophyll and hypocotyl cells. A. P5CS1-YFP localization in leaf mesophyll cells under control (−0.25 MPa) or low ψ_w_ stress (4 d at −0.7 MPa). Note that P5CS1-YFP level increased in the low ψ_w_ but P5CS1-YFP was preferentially localized close to chloroplasts in both treatments. Scale bars indicate 10 μm. B. P5CS1-YFP localization in hypocotyl epidermal cells under control (−0.25 MPa) or low ψ_w_ stress (4 d at −0.7 MPa). The preferential localization of P5CS1 around chloroplasts can be clearly seen in both treatments. Scale bars indicate 10 μm. C. Imaging of wild type hypocotyl cells using the microscope settings as in A and B demonstrated that there was no substantial bleed through of chloroplast autofluorescence into the YFP channel. Scale bars indicate 10 μm.

### High GT frequency and in-frame YFP insertion at the 3’ end of *AFL1*

We also tested GT insertion of YFP at the 3’ end of *AFL1*. The design of the DNA construct and sgRNA (Fig. 5A) following the methodology described above for *P5CS1*. This construct was then transformed into the T_2_ generation of *EC1.1_pro_:hypaCAS9* line #7. A first round of screening 30 T_2_ pooled DNA extracts (each pool contained seedlings from one T_1_ Basta-resistant plant, approximately 30 seedlings collected for each pool) identified six positive T_2_ DNA pools (Fig. 5B; Table 1). When individual plants from three selected T_2_ lines were screened, a high frequency of plants homozygous for the *YFP* insertion were found, including ten out of 24 plants for line #26 and #11 (Fig. 5C; Table 1; Supplemental Fig. S9). This contrasted with the *P5CS1* results and indicated that heritable homologous recombination occurred at both genomic copies of *AFL1* without lethality or disrupted development. Sequencing of genomic DNA from homozygous plants of line #26 confirmed that the donor fragment had been inserted intact with *YFP* in frame with the *AFL1* coding region (Supplemental Fig. S10).

**Figure 5:**
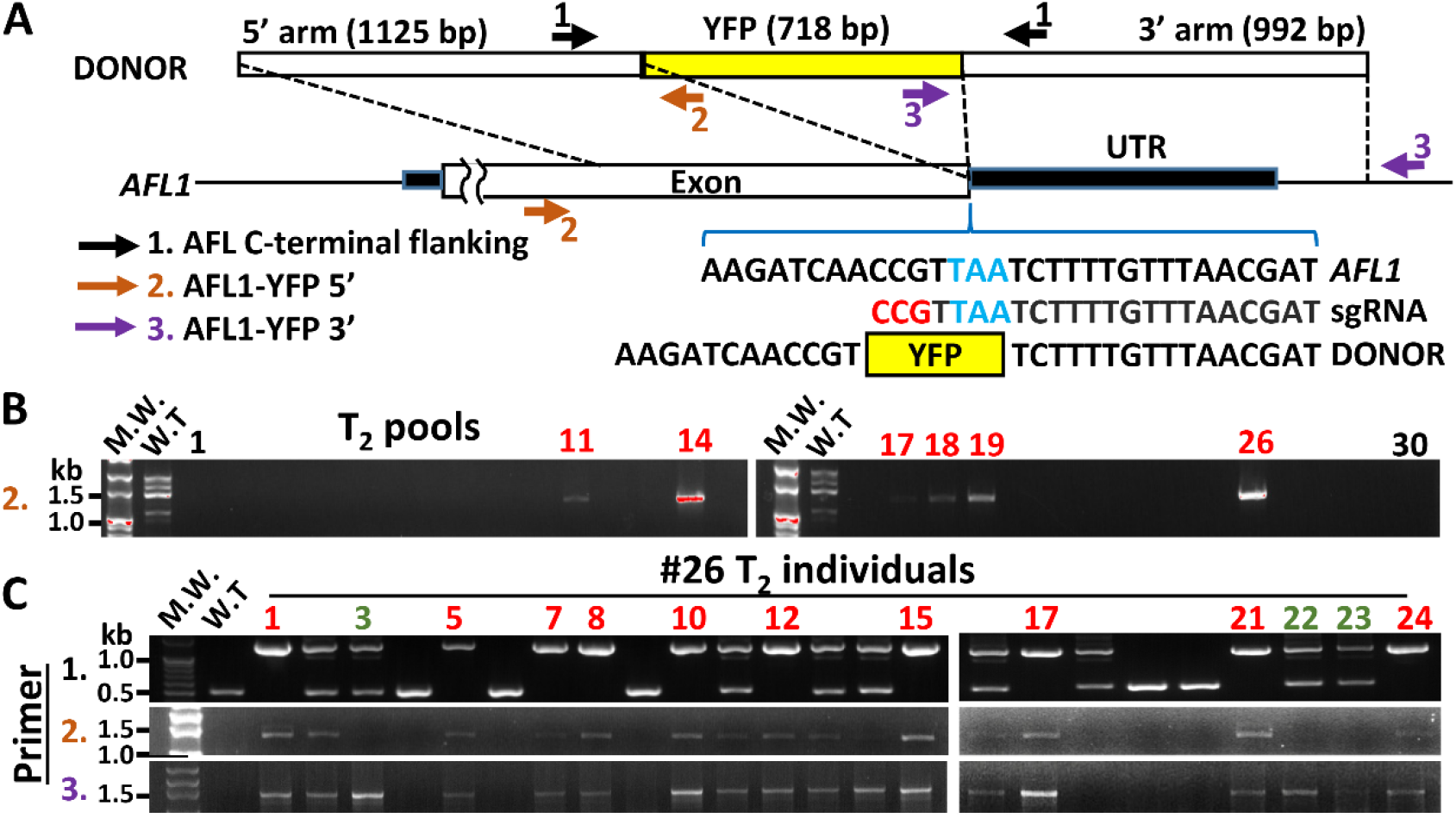
GT insertion of *YFP* at the 3’ end of *AFL1*. A. Donor Fragment design and genotyping primer positions for knock-in of C-terminal YFP at the *AFL1* locus. The *YFP* coding sequence was flanked by homology arms matching the indicated regions of *AFL1*. In the DNA sequences shown, blue indicates the AFL1 stop codon and red indicates the PAM site on the sgRNA. Note that replacement of *AFL1* endogenous sequence with the knock-in fragment disrupts the sgRNA recognition site thus preventing cutting of the donor fragment or re-cutting of endogenous *AFL1* after insertion of the donor fragment. Positions of primers used for genotyping are also shown (primer sequences and amplicon sizes can be found in Supplemental Table I). B. Genotyping of pooled T_2_ seedlings from 30 Basta resistant T_1_ plants using primer set 2. Red numbers indicate T_2_ seed pools with insertion of the YFP donor construct at the *AFL1* locus. Note that in this experiment, the WT control had multiple bands, possibly due to template DNA contamination. This did not affect interpretation of the results and individual plant screening (C) gave a single band of the expected M.W. C. Genotyping of individual T_2_ plants from line 26. Red numbers indicate plants homozygous for the *AFL1-YFP* knock-in. Green numbers indicate plants having putative single crossover event where homologous recombination occurred at the 3’ side of the donor fragment (indicated by the expected size band amplified by primer set 3) but other putative insertion or deletion occurred on the 5’ side of the donor fragment (based on lack of band amplified by primer set 2).

Interestingly, in line #14 amplification of the 3’ end of *AFL1* (primer set 3) revealed both the expected size band for *AFL1-YFP* generated by homologous recombination as well as a larger band (Supplemental Fig. S9). In these putative “single crossover” lines, it appears that for one chromosome homologous recombination occurred at the 5’ end of the donor construct, while at the 3’ end NHEJ occurred at the end or close to the end of the 3’ homology arm. This resulted in duplication of a part of the homology arm sequence, leading to the larger sized PCR band. Similarly, all of the AFL1-YFP GT lines assayed had some plants where the expected size band was amplified on the 5’ or 3’ end (primer set 2 or 3), but there was no band amplified on the other side of the YFP insertion. In these cases, it is also possible that a repair process incorporated only part of the donor construct (or there was deletion of chromosomal DNA), such that the primer site(s) for primer set 2 or 3 were disrupted. These data indicate that double strand breaks could be repaired by a combination of homologous recombination and NHEJ.

Homozygous plants from *AFL1-YFP* line #26 were backcrossed to wild type and *AFL1-YFP* transcript and protein levels were compared to wild type at both the T_3_ (before backcrossing) and T_6_ (after backcross) generations. The *AFL1* transcript levels were similar for the knock-in line and wild type in the T_3_ and T_6_ generation and for both control and stress treatments (Fig. 6A). Immunoblotting using AFL1-specific antisera (Kumar et al., 2015), demonstrated that T_3_ knock-in lines produced the AFL1-YFP fusion protein (approximately 27-30 kD higher molecular weight than wild type AFL1), and had no detectable production of untagged AFL1 (Fig. 6B). This was further confirmed by use of YFP-specific antisera which detected a band at the same molecular weight as AFL1-YFP but did not detect free YFP (Fig. 6B). However, in both the T_3_ and T_6_ generation, the level of AFL1-YFP protein detected was only about half the amount of untagged AFL1 detected in wild type (Fig. 6B, C and D). AFL1-YFP was detected in the hypocotyl under both control and low ψ_w_ treatments and was strongly induced by low ψ_w_ in young leaves (Supplemental Fig. S11A). Higher magnification images showed that AFL1-YFP was diffusely localized in the cytoplasm (Fig. S11B) and did not form foci or actin-associated strands as we previously observed in plants expressing *35S:YFP-AFL1* (Kumar et al., 2019; Kumar et al., 2015). This, along with the reduced level of AFL1-YFP despite wild type level of *AFL1* expression, suggests that the C-terminal YFP interferes with AFL1 localization and may decrease AFL1 stability. Together these results show that we could use GT for high-frequency insertion of YFP at the 3’ end of AHL1; however, the C-terminal YFP may alter AFL1 function and protein level for reasons unrelated to the GT procedure.

**Figure 6:**
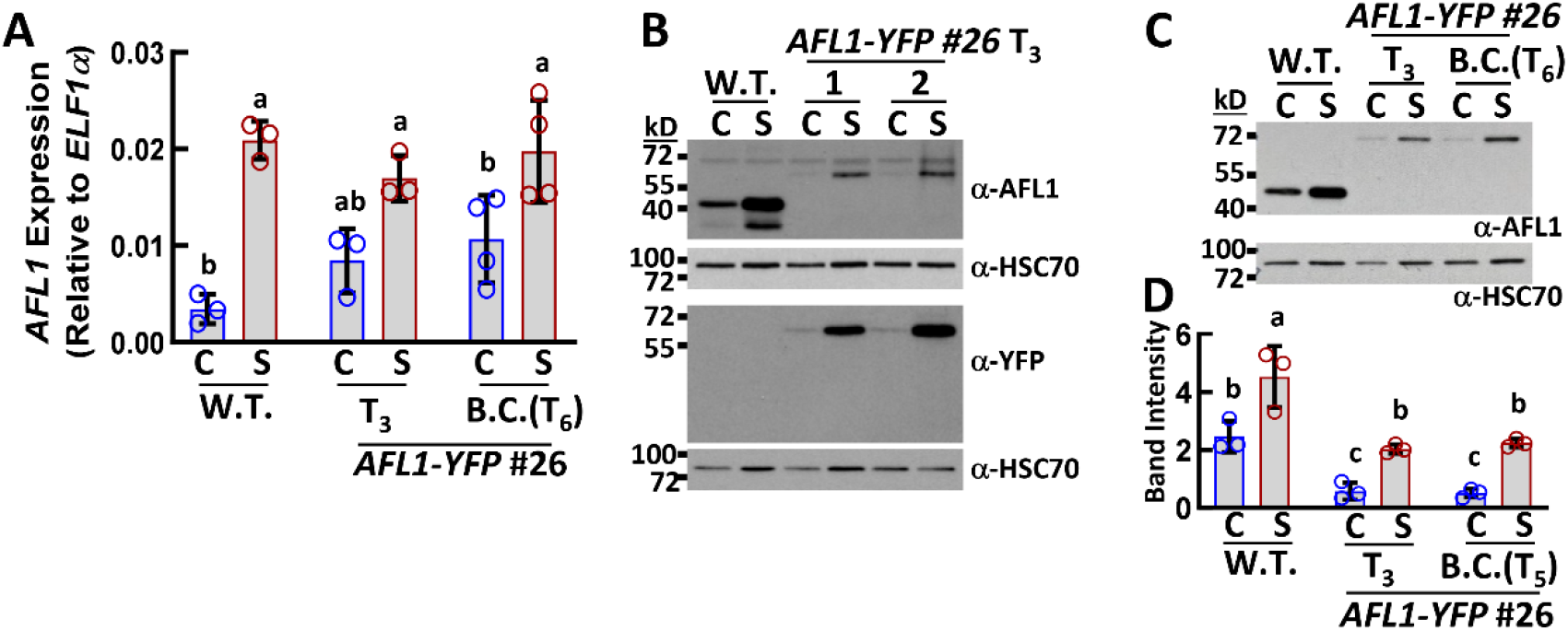
AFL1-YFP gene expression matches that of wild type but protein levels are reduced. A. QPCR of *AFL1* in wild type compared to T_3_ and T_6_ homozygous plants of *AFL1-YFP* knock-in line #26. Whole seedlings were collected at four days after transfer of seven-day-old seedlings to control or stress (−0.7 MPa) treatments (C = Control, S = Stress). Three or four biological replicates were performed. Data are means ±S.D. Data sharing the same letter are not significantly difference based on ANOVA with Tukey’s multiple comparison test (adjusted p ≤ 0.05). B. Immunoblot using antisera recognizing AFL1 (top) or YFP (bottom). In both cases, blots were stripped and re-probed with antisera recognizing HSC70 as a loading control. The AFL1 and YFP blots were prepared using the same samples. Protein was extracted from whole seedlings of wild type or T_3_ homozygous plants collected four days after transfer of seven-day-old seedlings to control or stress (−0.7 MPa) treatments (C = Control, S = Stress). 10 μg of protein was loaded in each lane. The predicted molecular weight of AFL1 is 42 kD. The predicted molecular weight of the AFL1-YFP fusion protein is 66 kD. C. Immunoblotting of T_6_ seedlings after backcrossing to wild type to remove CAS9 and the recombination donor construct. Experimental conditions were as described for E. D. Quantification of relative AFL band intensity in three biological replicates. Data sharing the same letter are not significantly different based on ANOVA with Tukey’s multiple comparison test (adjusted p ≤ 0.05).

### In-frame insertion of *YFP* at the 5’ end of *P5CS1* or *AFL1* disrupts expression

Given that for some proteins, such as AFL1, insertion of a tag sequence at the N-terminus rather than C-terminus is required, we also used our GT procedure to attempt such an insertion for *P5CS1* and *AFL1*. In this case we transformed the constructs containing donor DNA fragment and sgRNA into the T_2_ (for *AFL1*) or T_3_ (for *P5CS1*) generation of line #7 *EC1.1_pro_:hypaCAS9* plants. For *P5CS1*, two positive T_2_ seed pools were obtained (Supplemental Fig. S12A and B; Table 1). Genotyping of individual T_2_ plants showed that all plants of line #6 produced bands of the expected size at both 5’ and 3’ ends, and several homozygous plants were identified (Supplemental Fig. S12C). However, line #13 had a higher-than-expected molecular weight band at the 5’ end (primer set 3; Supplemental Fig. S12B, S13A). This was similar to *AFL1* C-terminal knock-in line #14 and indicated that the 3’ end of the donor construct had undergone NHEJ leading to duplication of the 5’ end of *P5CS1* homology arm.

For the *YFP*-*P5CS1* line #6, sequencing of genomic DNA from homozygous plants demonstrated that *YFP* was inserted intact and in-frame with *P5CS1* (Supplemental Fig. S13B). However, we found that the 5’knock-in lines had greatly reduced *P5CS1* transcript levels (Supplemental Fig. S12D) and neither YFP-P5CS1 nor untagged P5CS1 protein could be detected (Supplemental Fig. S12E). Consistent with this lack of P5CS1, low ψ_w_-induced proline accumulation was reduced to the same level as the *p5cs1-4* knock-out mutant (compare Supplemental Fig. S12F to Fig 2C). This absence of *P5CS1* gene expression, P5CS1 protein accumulation and proline accumulation was not rectified by backcrossing and advancing to the T_5_ generation (Supplemental Fig S12D and F). Thus, GT insertion of *YFP* at 5’ end of *P5CS1* abolished or strongly reduced *P5CS1* expression. However, the lack of homozygous lethality in the T_3_ generation indicated that the underlying mechanism was different than the situation observed for knock-in at the 3’ end of *P5CS1*.

Simultaneously with those experiments, we used a similarly designed construct for GT to insert *YFP* at the 5’ end of *AFL1* (Supplemental Fig. 14A). We identified several homozygous plants when screening individual plants from the positive T_2_ plants (Supplemental Fig. 14B and C). We also confirmed that genomic DNA of homozygous knock-in plants had an in-frame insertion of YFP at the AFL1 N-terminus (Supplemental Fig. S15). However, *AFL1* transcript levels were no longer induced by low ψ_w_ in knock-in lines (Supplemental Fig. 14D) and neither YFP-AFL1 nor untagged AFL1 could be detected in immunoblots using backcrossed T_6_ lines (Supplemental Fig. 14E). Thus, *P5CS1* and *AFL1* were similar in that GT insertion of *YFP* at the 5’ end of each gene strongly reduced, or even abolished, expression.

## Discussion

GT is a viable option for introducing many types of modifications to plant genomes. Our data provide an example of how relatively high frequency GT can be achieved in Arabidopsis even with a straightforward protocol. Additional refinements in terms of the type of *CAS* gene used, for example intronized *CAS9* (Grutzner et al., 2021) or *LbCas12a* (Merker et al., 2020; Wolter and Puchta, 2019), or refinements in the composition of the donor construct and the way in which the donor construct is delivered are likely to further increase GT frequency. As GT becomes more feasible, particularly in model plants, the next step in applying it for research is to better understand when it would be advantageous compared to the more traditional approach of transgenic complementation of a mutant with constructs encoding the protein of interest fused to a tag and expressed under control of its native promoter. To explore this question, we chose *P5CS1* and *AFL1*, in part because they have been difficult to study by other means. For *P5CS1*, it is unclear whether its promoter works properly in transgenic lines where it is put into a different genomic context and likely does not have its normal level of DNA methylation. For *AFL1*, the presence of transposable elements in the promoter region and 5’ UTR make it difficult to choose a promoter fragment to use in transgenic plant production. Our overall objective was to determine whether we could use GT to tag these two interesting stress-related proteins and thus generate tools for further study of protein localization and interaction. For that objective, the question of whether GT itself may alter expression of the target locus was of central importance.

We were ultimately successful in producing plants with *P5CS1-YFP* expression and protein accumulation at the same level as wild type *P5CS1* and having the same level of stress-induced proline accumulation. We also readily identified homozygous plants where *AFL1-YFP* had the essentially the same level of gene expression as wild type *AFL1*. At the same time, our experiments made it clear that GT itself can alter expression of the target gene and this potential complication should be monitored when using GT for functional analyses. Homozygous *P5CS1-YFP* plants with wild type levels of expression could only be found in the T_5_ and later generations, possibly because of epigenetic disruption that took several generations to reset to the wild type state. Also, insertion of YPF at the 5’ end of *AFL1* or *P5CS1* (to produce *YFP-AFL1* and *YFP-P5CS1*) greatly reduced, or abolished the expression of either gene. In contrast to the *P5CS1-YFP* results, backcrossing and advancement to the T_6_ generation did not lead to recovery of *YFP-AFL1* or *YFP-P5CS1* expression. The reasons for this are unknown. One possibility is that the linker sequence inserted between *YFP* and the *AFL1* or *P5CS1* coding region disrupted expression. However, this is unlikely as we used the same linker sequence present in various Gateway cloning plant transformation vectors in which high levels of N-terminal YFP fusion protein are produced. Also, the linker sequence did not interfere with expression of *P5CS1-YFP* or *AFL1-YFP*.

An obvious caveat to these observations is that two genes is a small number from which to make any broad conclusions. Moreover, the initial lethality of *P5CS1-YFP* insertion was only observed for one gene. As stated above, *P5CS1* and *AFL1* were selected for these experiments because of factors (promoter methylation, nearby transposons) that made them problematic for traditional transgenic approaches. Thus, it is possible that these genes are also especially problematic for GT. Because of this, we do not offer any prediction as to how often the problems we encountered will come up, merely that they can. At the same time, we note that this caveat of having only examined a few example genes also applies to other GT studies, many of which have used a relatively limited number of easy-to-assay loci or have focused on insertion into a limited number of “safe” intergenic sites where a whole construct can be added without disrupting any endogenous genes. There is at least one report that GT-mediated insertion of RFP was found to alter DNA methylation status (Lieberman-Lazarovich et al., 2013). Conversely, GT frequency can be dramatically affected by chromatin remodeling (Shaked et al., 2005). Thus, GT disruption of the heavily methylated regions close to P5CS1 or disruption of the epigenetic status of transposable elements in the promoter and 5’ UTR of AFL1may have been a factor in how GT disrupted expression of these genes. It has also been observed that GT frequency is affected by temperature or light conditions (Vu et al., 2020), and more efficient GT was observed after heat shock to induce CAS9 expression (Barone et al., 2020), both of which may also be related to changes in chromatin status. Despite the high interest in GT, we are not aware of any established guidelines to indicate which type of gene (i.e., what type of epigenetic landscape) is likely to allow for high frequency GT, or conversely, which sites are likely to be more recalcitrant to GT and (or) prone to exhibit altered gene expression after GT. Tests of GT at additional loci differing in DNA methylation or chromatin state will be useful to determine whether our experience with *P5CS1* and *AFL1* is an outlier or will be encountered more frequently.

The *P5CS-YFP* knock-in line generated here has already allowed us to address long standing questions about P5CS1 tissue-specific accumulation and subcellular localization. Szekely et al. (2007) expressed P5CS1-GFP under control of 2426 bp of endogenous promoter sequence and observed a low level of protein accumulation in unstressed shoot tissue and in mature roots. In contrast to our results, they did not observe P5CS1-YFP accumulation in root tips and did not report accumulation of P5CS1 in guard cells. The guard cell accumulation of P5CS1 is consistent with previous transcript profiling of guard cells (Yang et al., 2008) and shows that proline metabolism in guard cells should receive more attention in terms of how it affects stomatal function. High levels of P5CS1-YFP in the root meristem and elongating cells is consistent with observations that *p5cs* mutants have impaired root growth, perhaps by affecting reactive oxygen levels in the root meristem (Biancucci et al., 2015; Bauduin et al., 2022). Also, our observation that P5CS1 highly accumulates in root steele cells, but not in other root cell types agrees with previous transcriptional profiling (Dinneny et al., 2008) and suggests that proline accumulation is less cell-autonomous than previously thought. Our observation that P5CS1 accumulates in the growing tip of root hairs is also surprising and it will be of interest to test whether tip growing cells such as root hairs and trichomes are altered in proline metabolism mutants.

In terms of P5CS1 subcellular localization, Szekely et al. (2007) reported that P5CS1 localized in cytoplasmic foci. This contrasted with P5CS2-GFP, which was diffusely localized in cytoplasm. They also concluded that salt or osmotic stress changed P5CS1 localization from cytoplasm to chloroplast. Our observations do not support this later conclusion but rather show that P5CS1 clusters around the outside of chloroplasts. These observations fit with observations that disruption of P5CS1 changes photosynthesis-related gene expression (Shinde et al., 2016), and that light has a strong and direct influence on *P5CS1* expression and proline accumulation (Abraham et al., 2003; Feng et al., 2016; Kovacs et al., 2019). Consistent with our results, Funck et al. (2020) did not observe chloroplast-localized P5CS1. However, they only analyzed protoplasts isolated from plants expressing P5CS1-YFP under control of an ubiquitin promoter and did not examine the effects of stress on P5CS1 localization. Our P5CS1-YFP line will be useful for further studies to examine P5CS1 interactions and post-translational modifications. Such data can reveal the mechanisms of how proline is linked to cellular redox status and to other specific metabolic pathways such as the oxidative pentose phosphate pathway (Zheng et al., 2021). Because we anticipate the P5CS1-YFP line to have stable expression pattern, and because it no longer has a selectable marker, many strategies such as crossing this line to other mutants, transforming it, or using it as the basis for forward genetic screening, are now feasible. Overall, our results both show how GT can a basis for analysis of gene function and generate a stable P5CS1-YFP line that will be a unique resource for study of one of the most prominent abiotic stress-related genes.

## Author Contributions

T.L. and P.E.V. conceived the research; T.L., L.G. and P.E.V. designed the experiments; L.G. supplied key materials, T.L. and P.E.V. analyzed the data; T.L., H.-Y.C, T.C.L., C.-Y.C. and H.P. performed the experiments; P.E.V. wrote the article with assistance from T.L.; W.S. provided material, and W.S., T.L. and L.G. edited the article.

## Acknowledgements

We thank the staff of the imaging core facility of the Institute of Plant and Microbial Biology (Ji-Ying Huang and Mei-Jane Fang) for microscopy assistance and Shih-Shan Huang for laboratory assistance. This research was supported by Academia Sinica Investigator Awards to P.E.V (AS-IA-108-L04) and W.S. (AS-IA-111-L03).

## Conflict of Interest Statement

The authors declare that they have no conflict of interests.

## Notes

### Competing Interest Statement

The authors have declared no competing interest.

### Summary of Updates

The paper has been updated to incorporate new data on the successful generation of P5CS1-YFP knock in plants and new information on P5CS1 expression pattern and subcellular localization.

