## Supplemental Figures S1-S15 for "Insertion of *YFP* at *P5CS1* and *AFL1* shows the potential, and potential complications, of gene tagging for functional analyses of stress-related proteins"

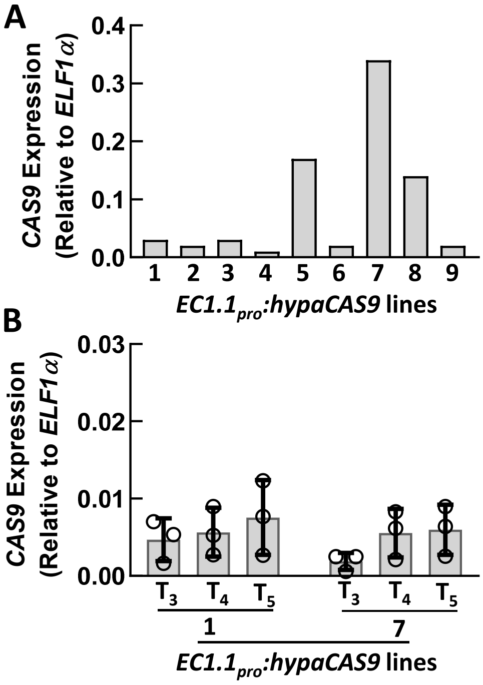


**Figure S1: *HypaCAS9* gene expression in transgenic lines.**

A. Expression of *HypaCAS9* in $T_2$ transgenic lines. One sample from each line was assayed to give an estimate of *HypaCAS9* gene expression while still preserving other plants for retransformation. Floral buds were sampled from one plant for each line. Note that in these initial tests only a single sample from each line was used for QPCR assays as we wanted to preserve as much of the $T_2$ seed as possible for the second stage transformation.

B. Expression of *HypaCAS9* in $T_3$ and later generations of plants derived from two $T_2$ lines with contrasting expression. Three biological replicates were assayed, each containing floral buds from one or two plants. Data are means ± S.D. (n = 3).

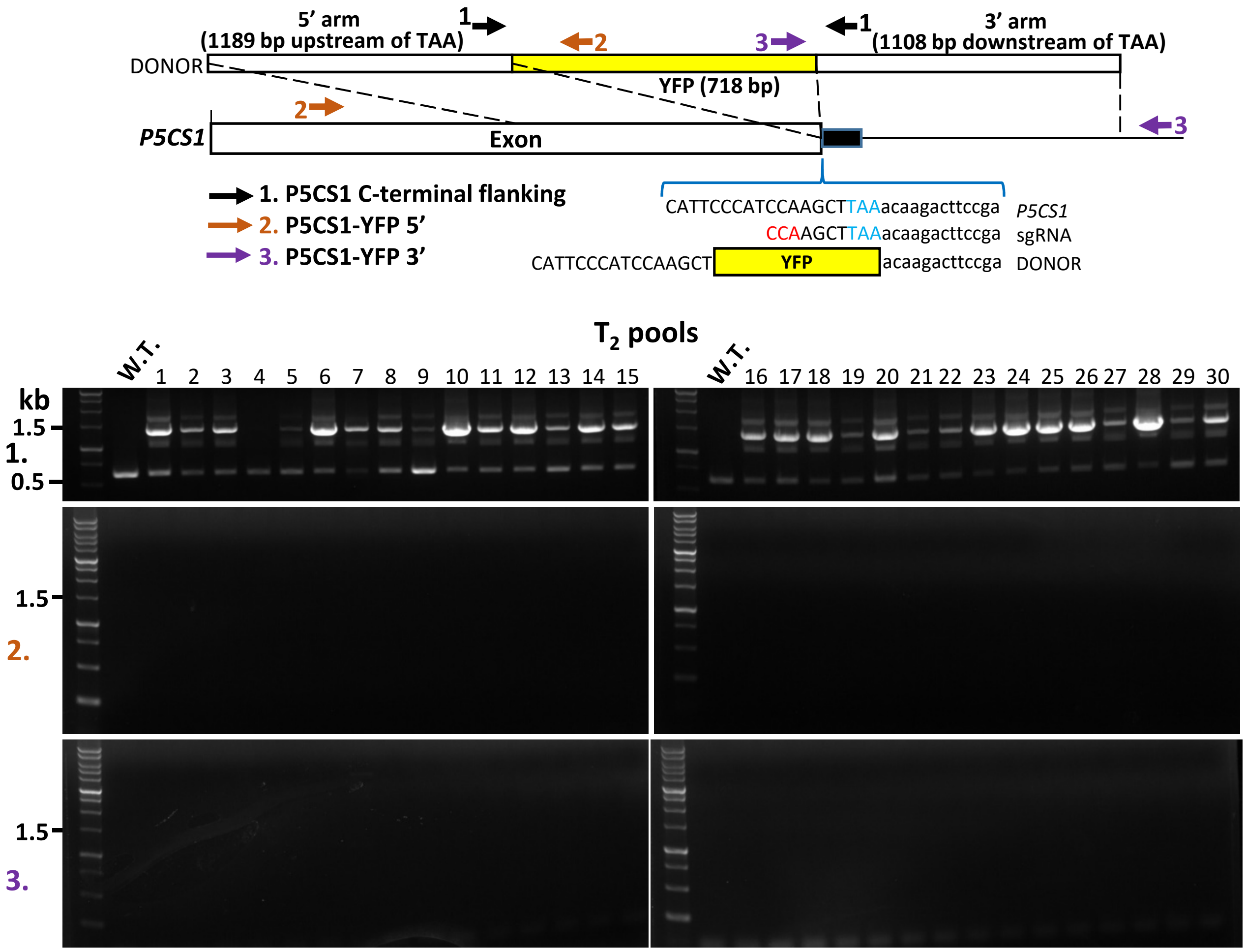


**Figure S2: Knock-in construct and screening for *P5CS1* 3' (*P5CS1-YFP*) GT using $T_3$ generation of CAS9 line #7.**

Top part shows the Donor Fragment design and genotyping primer positions for knock-in of YFP at the 3' end of *P5CS1*. The diagram is repeated here from Fig 1A for clarity of interpreting panel B. The YFP coding sequence was flanked by homology arms matching the indicated regions of *P5CS1*. In the DNA sequences shown, blue indicates the *P5CS1* stop codon and red indicates the PAM site on the sgRNA. Note that replacement of *P5CS1* endogenous sequence with the knock-in fragment disrupts the sgRNA recognition site. Positions of primers used for genotyping are also shown (primer sequences and amplicon sizes can be found in Supplemental Table I).

Bottom part shows genotyping of $T_2$ seed pools from transformation of the $T_3$ generation of CAS9 starter line #7. In the $T_2$ generation after retransformation nearly all seed pools tested had the donor construct T-DNA integrated into genomic DNA (as indicated by the high M.W. band amplified by primer set 1), but none had insertion of *YFP* at the *P5CS1* as indicated by the lack of amplification for primer sets 2 and 3.

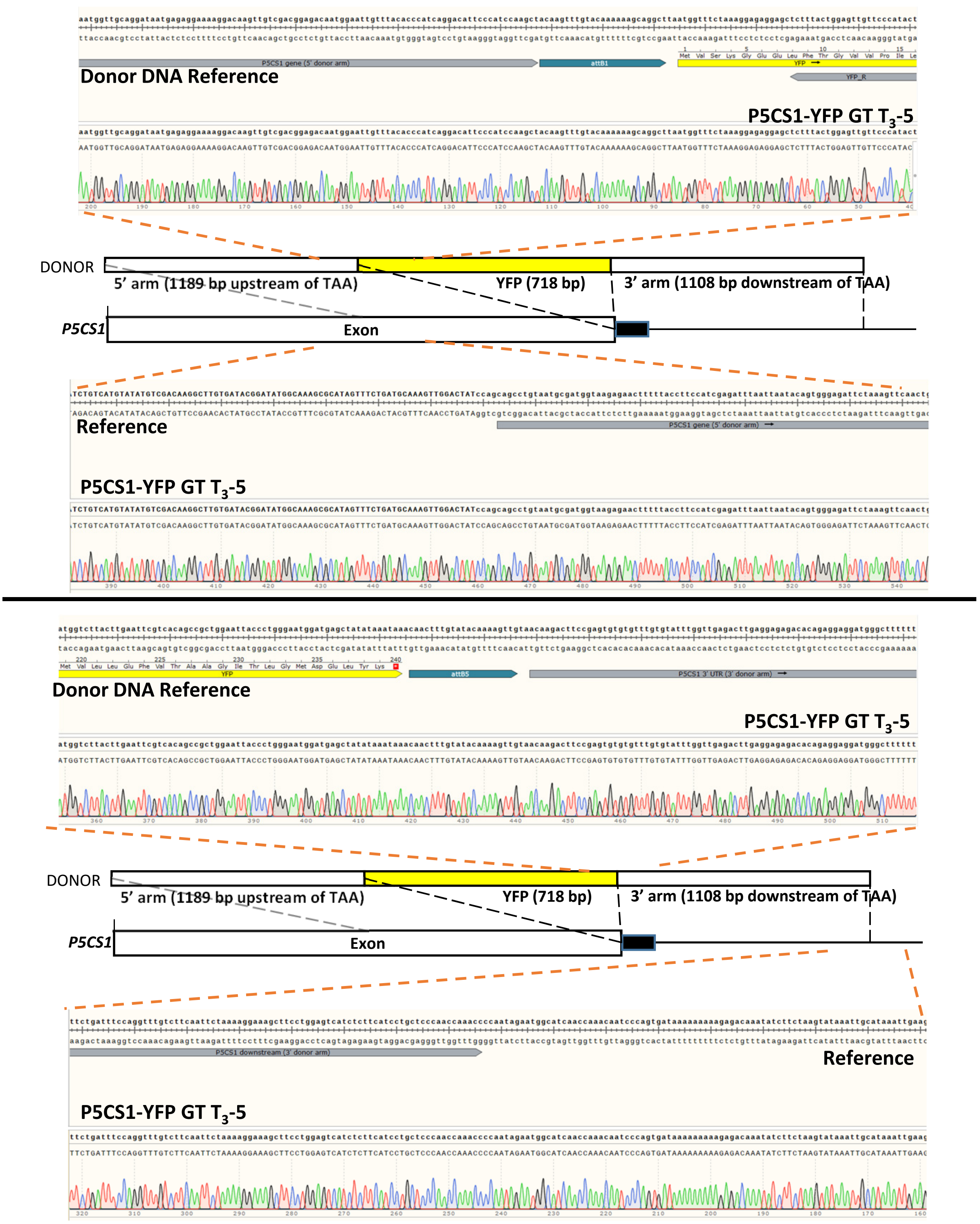


**Figure S3:** Sequencing through the ends of the homology arms and the YFP fusion to P5CS1 coding region and 3' UTR confirms that the GT insertion occurred via homologous recombination and did not alter P5CS1 coding region or UTR sequence at the junction of the homology arm and the endogenous gene and the YFP coding sequence remained in-frame with *P5CS1*.

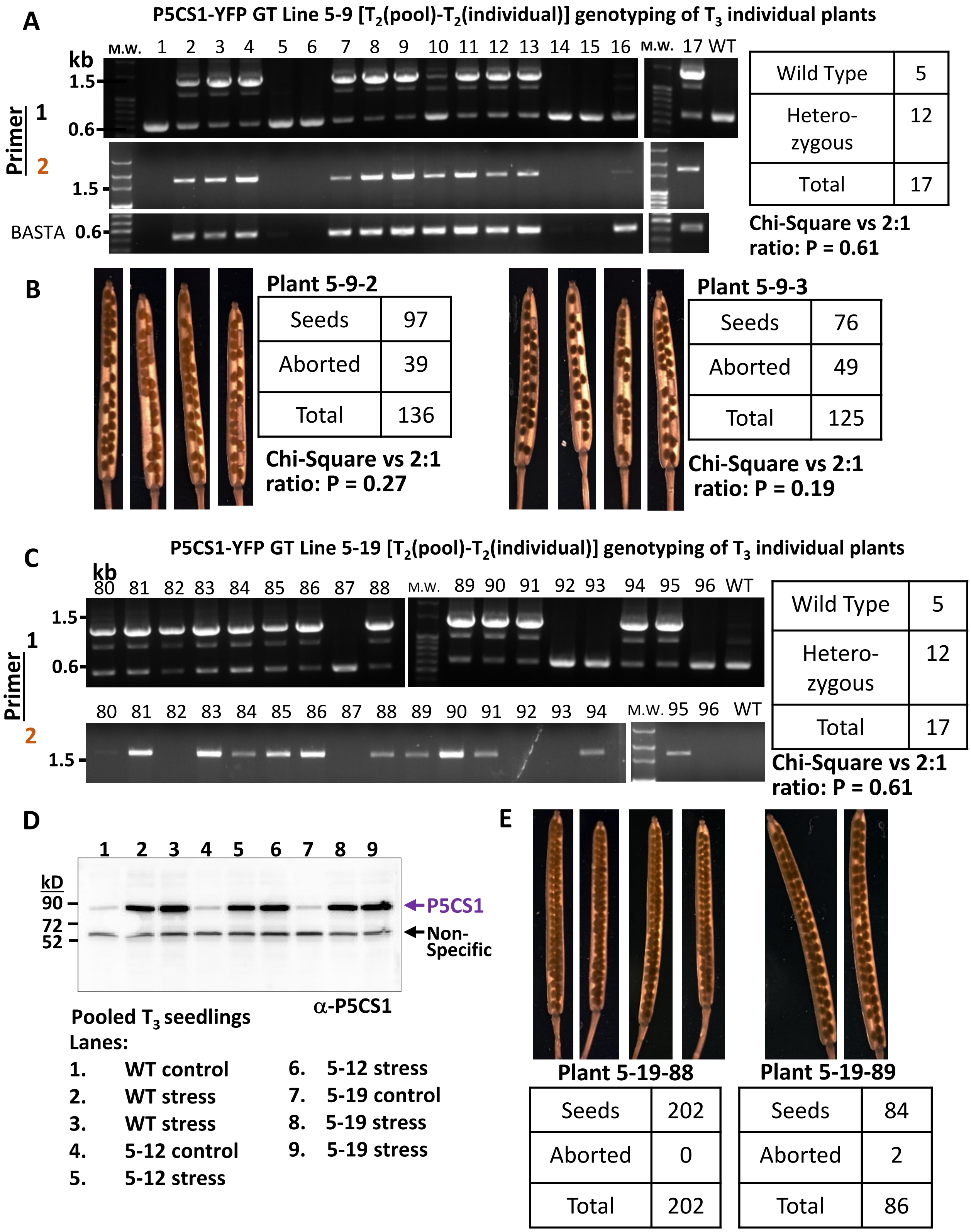


**Figure S4: $T_3$ genotyping of P5CS1-YFP GT lines. (Legend on next page)**

**Figure S4: $T_3$ genotyping of P5CS1-YFP GT lines.**

A. Genotyping individual $T_3$ plants from line 5-9. The result is representative several $T_3$ lines genotyped. Primer 1 and 2 are shown in Fig. 1A and Fig S2. Primer 1 amplifies the *P5CS1* sites flanking the site of YFP insertion and will amplify a larger size band from the donor DNA fragment or from the *P5CS1* gene after *YFP* insertion but 1 amplify a shorter fragment from the wild type *P5CS1* allele. Thus, presence of the shorter wild type band indicates that the plants are wild type or heterozygous for the *YFP* insertion. Primer 2 amplifies from *P5CS1* outside the homology are of the donor DNA and from within the YFP sequence and thus can only amplify a fragment when *YFP* has been inserted into the endogenous DNA. Primers specific to the BASTA resistance gene were used to confirm presence or absence to the T-DNA construct containing the donor DNA. Segregation ratio of heterozygous versus wild type plants was tested by Chi-square analysis vs the 2:1 ratio expected if homozygous *P5CS1-YFP* is lethal and thus not present in the seedlings analyzed.

B. Images of representative siliques containing $T_4$ seed from line 5-9-2 and 5-9-3 along with counts of aborted seeds (empty spaces in the silique) versus present seed and Chi-Square tests. Similar results were seen for other $T_3$ plants.

C. Genotyping individual T3 plants from line 5-19. Data presentation and analysis are as described for A except that the BASTA specific primer was not used in this case.

D. Immunoblot of P5CS1 shows that two representative $T_3$ lines lacked detectable level of the P5CS1-YFP fusion protein despite being heterozygous for the P5CS1-YFP knock-in. Note that the band at approximately 55 kd is non-specific and serves as a loading control.

E. Siliques containing T4 seed and segregation ratio of present versus aborted seed for two plants from the 5-19 line.

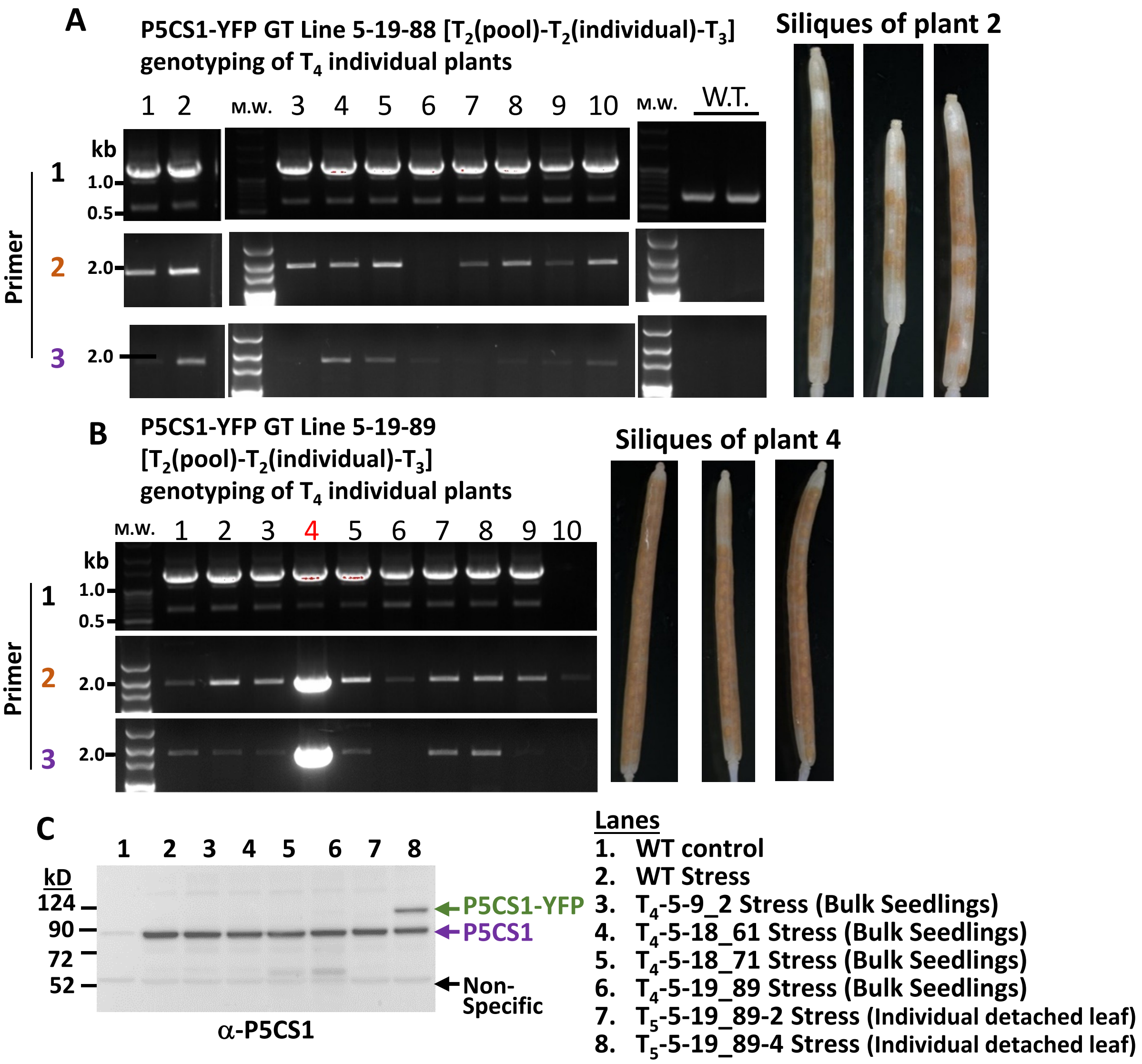


**Figure S5: $T_4$ genotyping of P5CS1-YFP GT lines.**

A. Genotyping individual $T_4$ plants from line 5-9-18. The result is representative several $T_4$ lines genotyped. Primer positions are as shown in Fig. 1A and Fig. S2. The lack of homozygous plants recovered and examination of representative siliques showed the continued high number of missing seed indicating that the P5CS1-YFP insertion continue to be homozygous lethal in this line.

B. Genotyping individual $T_4$ plants from line 5-9-18. Examination of siliques from plant 4 indicated that it had fewer missing seed than others. Also, screening of leaf samples for YFP fluorescence found positive result for plant 4 but not for others.

C. Immunoblot detection of P5CS1 in seedlings of the indicated lines or individual leaves of plant number 2 and 4 (leaves were detached and placed on -0.7 MPa PEG plates for 24 h to induce P5CS1 accumulation). Seedlings were stress treated by transfer to -0.7 MPa for four days as described for other experiments.

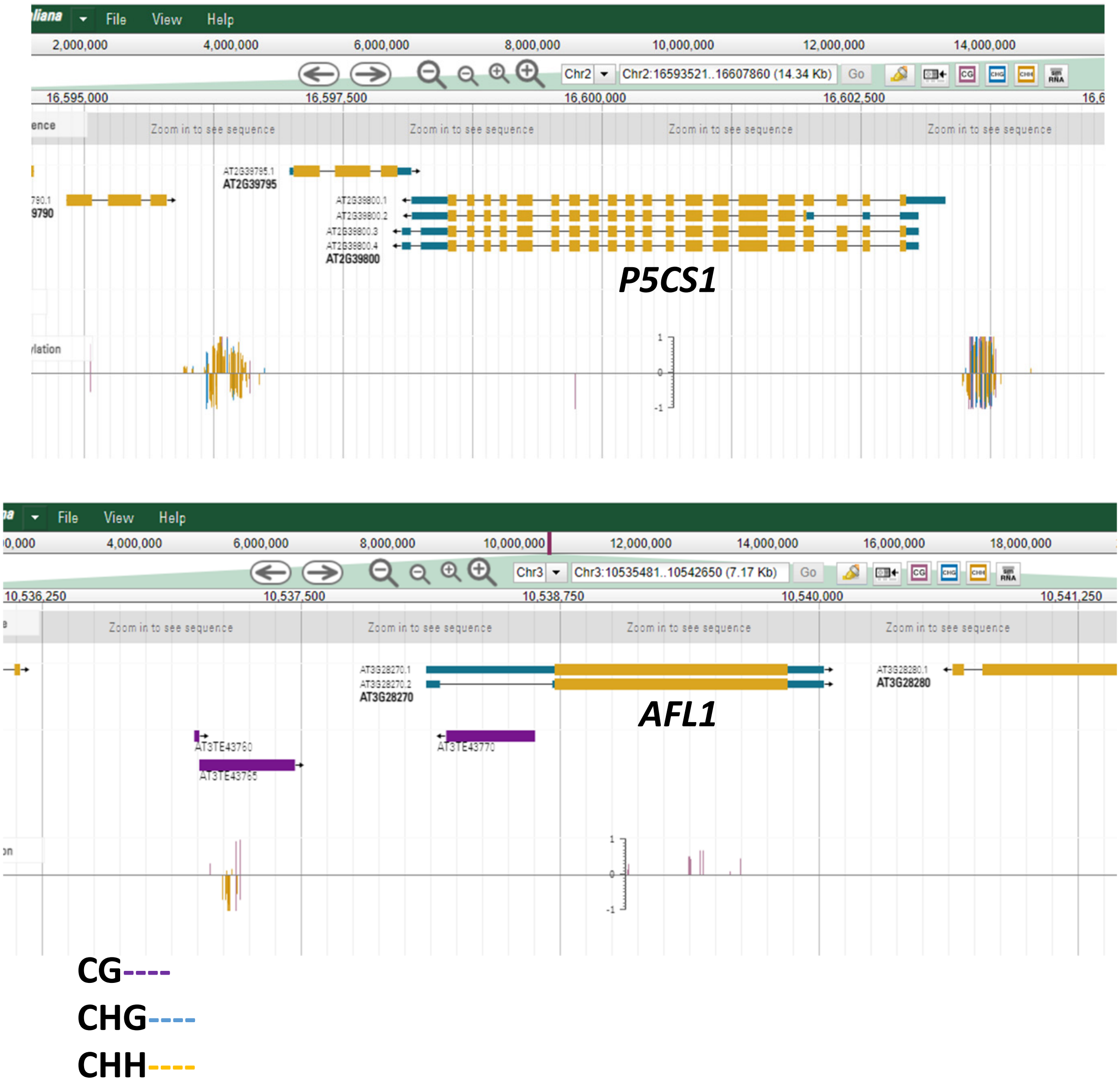


**Figure S6: Methylation patterns of *P5CS1* and *AFL1*.**
Data were retrieved from the Plant Methylome Database. Note the high level of methylation in both the *P5CS1* promoter and downstream of *P5CS1* in the promoter of AT2G39795. Also note the presence of transposable elements in the promoter and 3' UTR of AFL1.

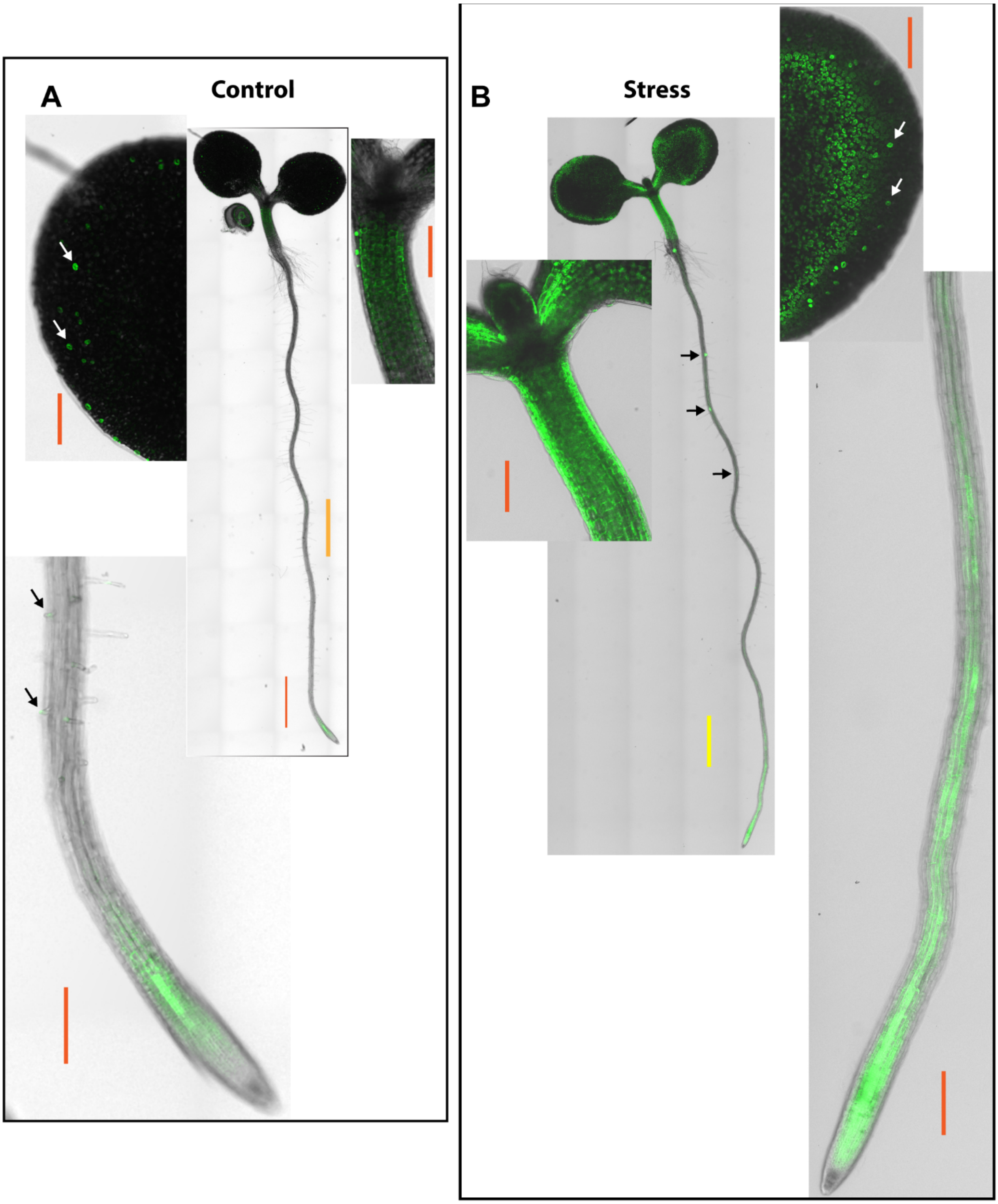


**Figure S7: Pattern of P5CS1-YFP accumulation under control and low $\psi_w$ stress (legend on next page).**

**Figure S7: Pattern of P5CS1-YFP accumulation under control and low $\psi_w$ stress.**

A. P5CS1-YFP accumulation in 3-day-old unstressed seedling. Center image shows whole seedling (yellow scale bar indicates 100 μm) while other images show enlarged images of shoot meristem, cotyledon and root (orange scale bars indicate 20 μm). In the cotyledon image, white arrows indicate P5CS1-YFP in guard cells. In the root image, black arrows indicate P5CS1-YFP in the tips of root hairs.

B. P5CS1-YFP accumulation in 4-day-old seedling that had been exposed to low $\psi_w$ stress (-0.7 MPa) for 24 h. Center image shows whole seedling (yellow scale bar indicates 100 μm) while other images show enlarged images of shoot meristem, cotyledon and root (orange scale bars indicate 20 μm). In the cotyledon image, white arrows indicate P5CS1-YFP in guard cells while the other signal is from mesophyll cells (see also Fig. 3A and Fig. S8B that shows lack of P5CS1-YFP in pavement cells). In the main image, black arrows indicate P5CS1-YFP in the tips of emerging lateral roots.

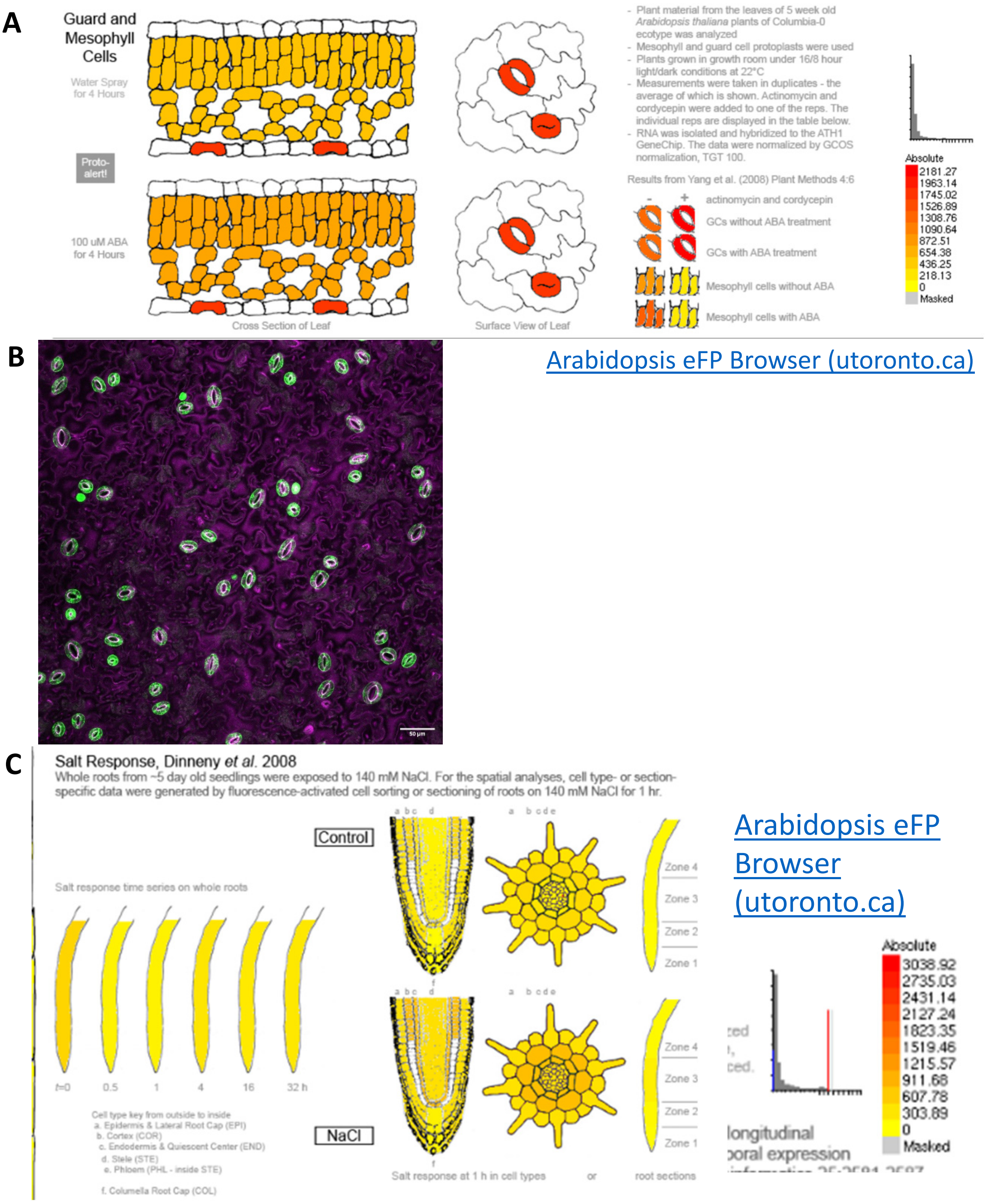


**Figure S8: *P5CS1* gene expression pattern from the eFP Browser.**

A. High constitutive expression of *P5CS1* in guard cells as well as ABA inducible expression in mesophyll cells.
B. Representative image of P5CS1-YFP in unstressed leaf of 11 day old seedling. The specific accumulation of P5CS1 in guard cells is consistent with the previously reported gene expression data.
C. Abiotic stress induced expression of *P5CS1* in cortex of the root tip is consistent with the patterns of P5CS1-YFP accumulation shown in Figure 3B.

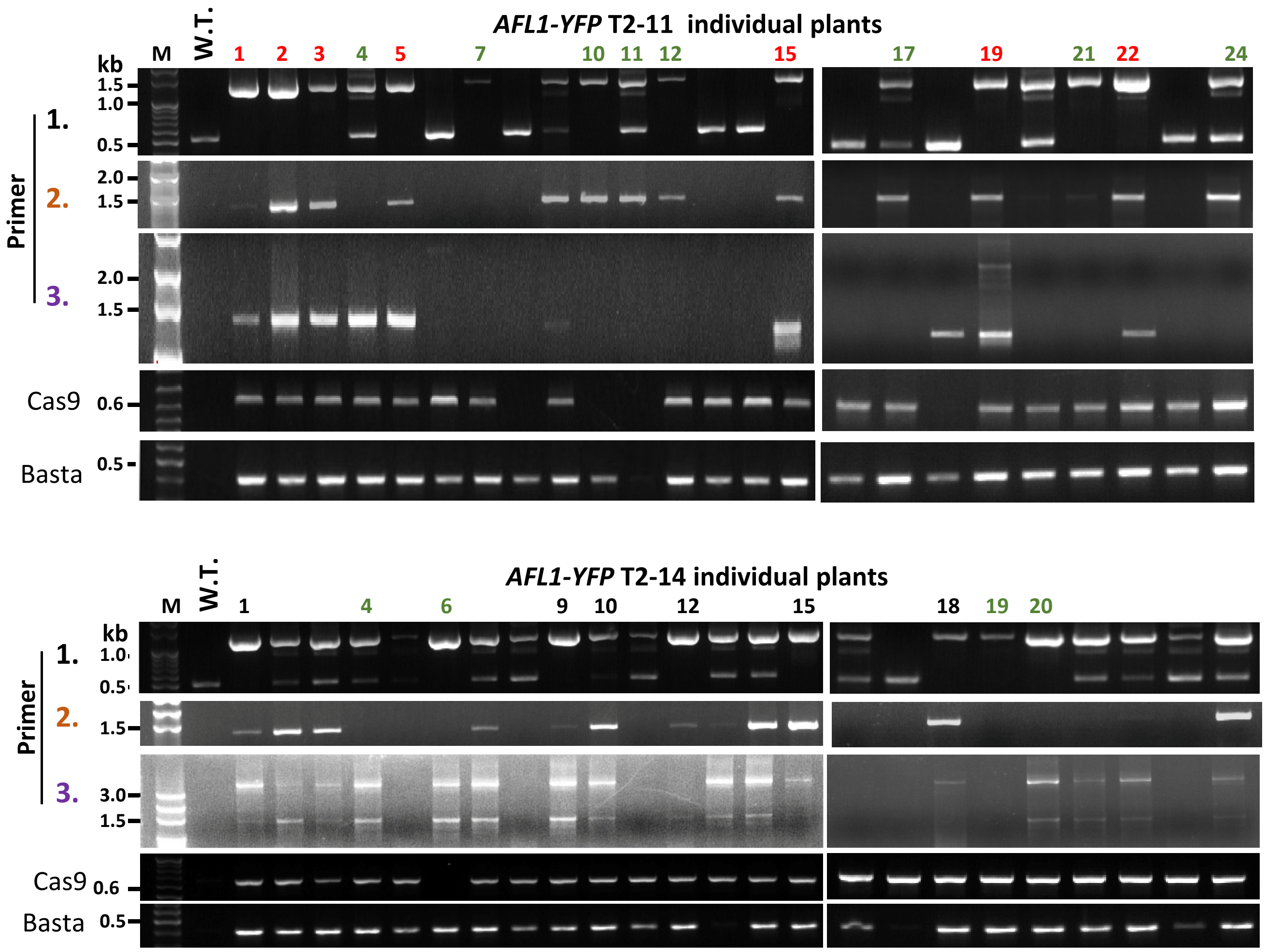


**Figure S9: PCR screening of additional $T_2$ lines of *AFL1* 3' knock-in.**
Primers and PCR conditions are as described for Figure 5. In addition, portions of the *Cas9* and *Basta* resistance genes were amplified to check whether CAS9 was still present and whether the construct from the second transformation of the recombination donor fragment had stably integrated into the genome.

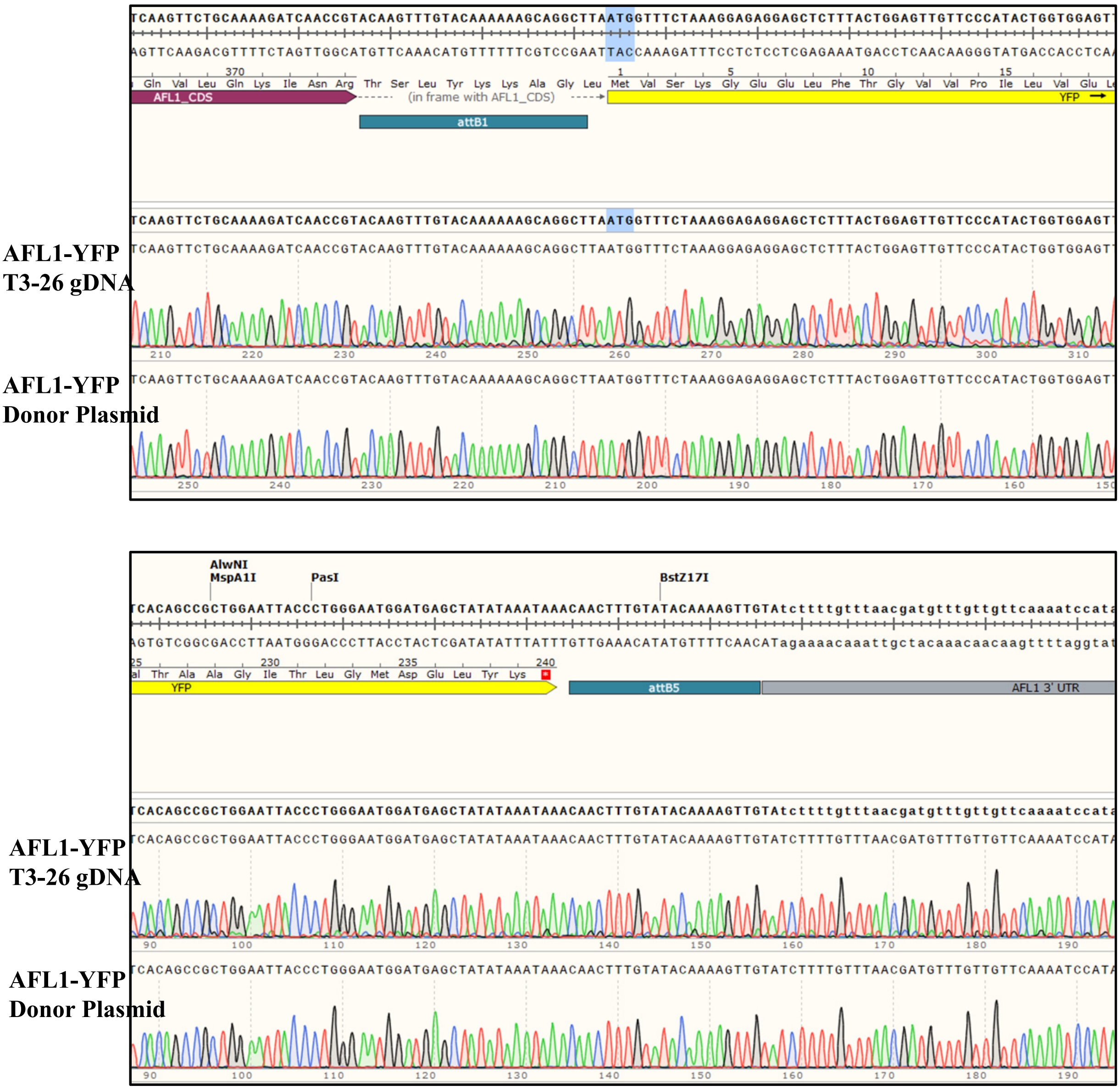


**Figure S10: Sequencing to show intact and in-frame insertion of YFP at the 3' end of *AFL1*.**

Sequencing chromatogram of *AFL1* 3' (*AFL1-YFP*) knock-in T3-26 genomic DNA (gDNA) aligned with the sequence of the donor fragment used in the second transformation. Alignments were conducted using SnapGene software. Results show that the both the 5' and 3' ends of the *YFP* sequence and flanking Gateway linker sequences used to prepare the construct were intact and integrated into genomic *AFL1*.

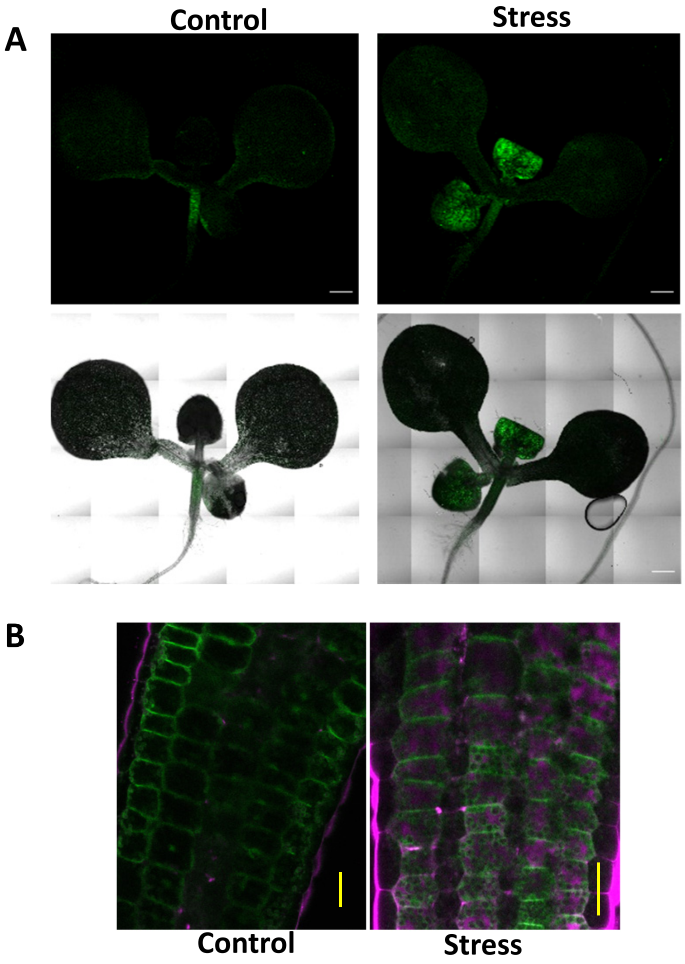


**Figure S11: Pattern of AFL1-YFP protein accumulation.**
A. AFL1-YFP protein accumulation pattern in $T_6$ seedlings under control or stress (four days after transfer to -0.7 MPa conditions. Scale bars indicate 500 μm.
B. Higher magnification images of AFL1-YFP in petiole cells from control or stress treated seedlings. Scale bars indicate 50 μm.

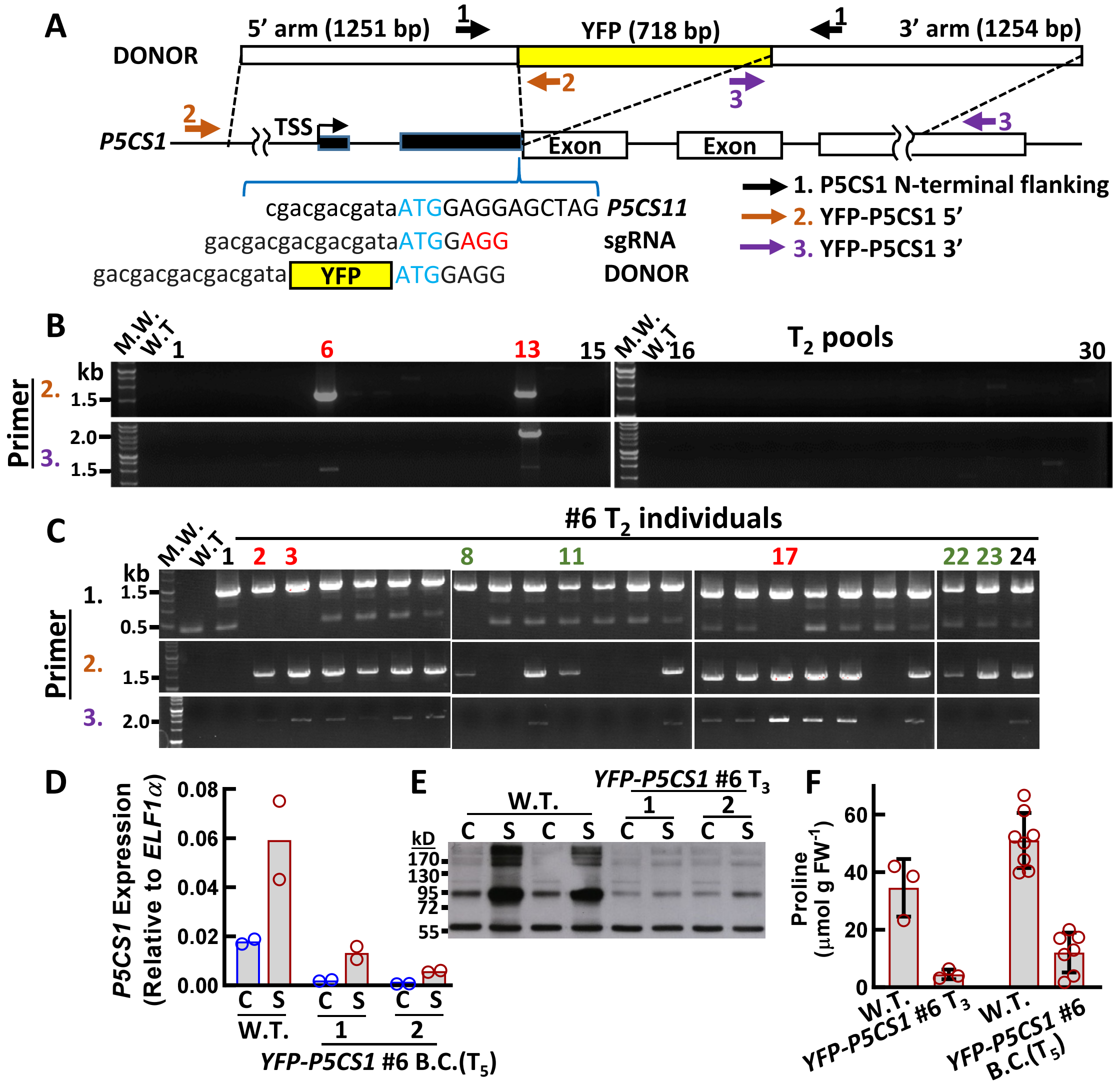


**Figure S12: N- terminal knock-in for *P5CS1* (legend on next page)**

**Figure S11: N- terminal knock-in for *P5CS1***

A. Donor Fragment design and genotyping primer positions for knock-in of N-terminal YFP at the *P5CS1* locus. The YFP coding sequence was flanked by homology arms matching the indicated regions of *P5CS1*. In the DNA sequences shown, blue indicates the *P5CS1* start codon and red indicates the PAM site on the sgRNA. Note that replacement of *P5CS1* endogenous sequence with the knock-in fragment disrupts the sgRNA recognition site. Positions of primers used for genotyping are also shown (primer sequences and amplicon sizes can be found in Supplemental Table I).

B. Genotyping of pooled $T_2$ seedlings from 30 Basta resistant $T_1$ plants using primer set 2. Red numbers indicate T2 seed pools with insertion of the YFP donor construct at the *P5CS11* locus.

C. Genotyping of individual T3 plants from line 26. Red numbers indicate plants homozygous for the *YFP-P5CS1* knock-in. Green numbers indicate plants having putative single crossover event where homologous recombination occurred at the 5' side of the donor fragment (indicated by the expected size band amplified by primer set 2) but other putative insertion or deletion occurred on the 3' side of the donor fragment (based on lack of band amplified by primer set 3).

D. QPCR of *P5CS1* in wild type compared to $T_3$ and $T_6$ homozygous plants of *YFP-P5CS1* knock-in line #26. Whole seedlings were collected at four days after transfer of seven-day-old seedlings to control or stress (-0.7 MPa) treatments (C = Control, S = Stress). Two biological replicates were performed.

E. Immunoblot using antisera recognizing P5CS1. Protein was extracted from whole seedlings of wild type or T2 homozygous plants collected four days after transfer of seven-day-old seedlings to control or stress (-0.7 MPa) treatments (C = Control, S = Stress). 10 mg of protein was loading in each lane. The predicted molecular weight of P5CS1 is 78 kD; however, we routinely observe that P5CS1 runs at higher than predicted molecular weight (Kesari et al., 2012). Note that the YFP-P5CS1 lanes, the faint bands seen in the 90-170 kD molecular weight range are consistent with previous observations of low level of cross reactivity of this antisera with P5CS2. The band at 57 kD is non-specific and serves as a loading control.

F. Proline accumulation in wild type and YFP-P5CS1 seedlings. Samples were collected four days after transfer of seven-day-old seedlings to control or stress (-0.7 MPa) treatments. Two or three independent experiments were performed and combined data of 3-8 samples from those experiments is shown. Error bars indicate S.D.

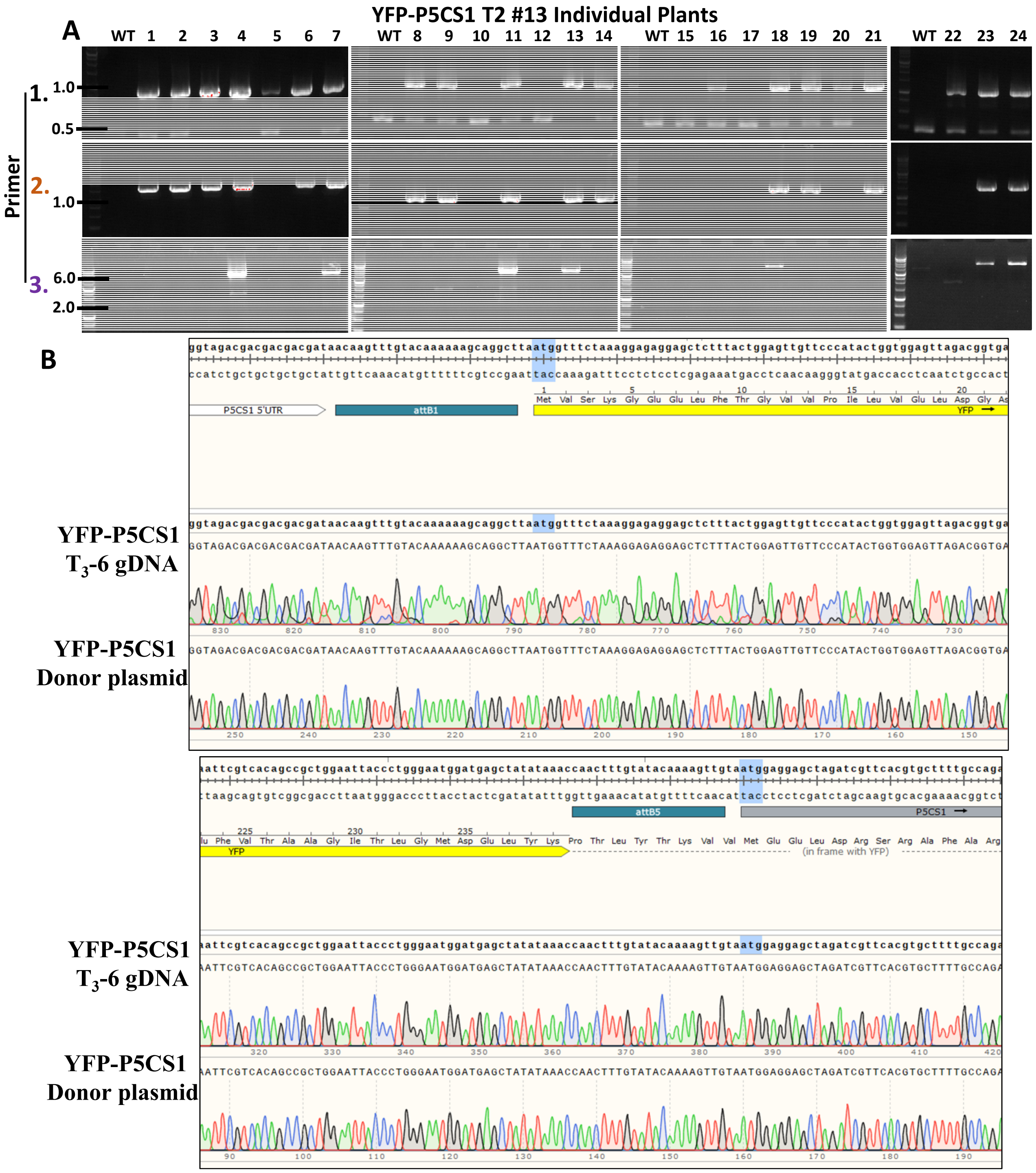


**Figure S13: Additional genotyping and sequencing to show the intact and in-frame insertion of *YFP* at the 5' end of *P5CS1*.**

A. Genotyping of individual plants from an additional *P5CS1* 5' (*YFP-P5CS1*) knock-in T2 line. Primers and PCR conditions are as described for Figure 4.

B. Sequencing chromatogram of *P5CS1* 5' knock-in T3-6 genomic DNA (gDNA) aligned with the sequence of the donor fragment used in the second transformation. Results show the successful in-frame integration of *YFP*, along with flanking Gateway recombination sequences used to prepare the donor construct, at the 5' end of *P5CS1*.

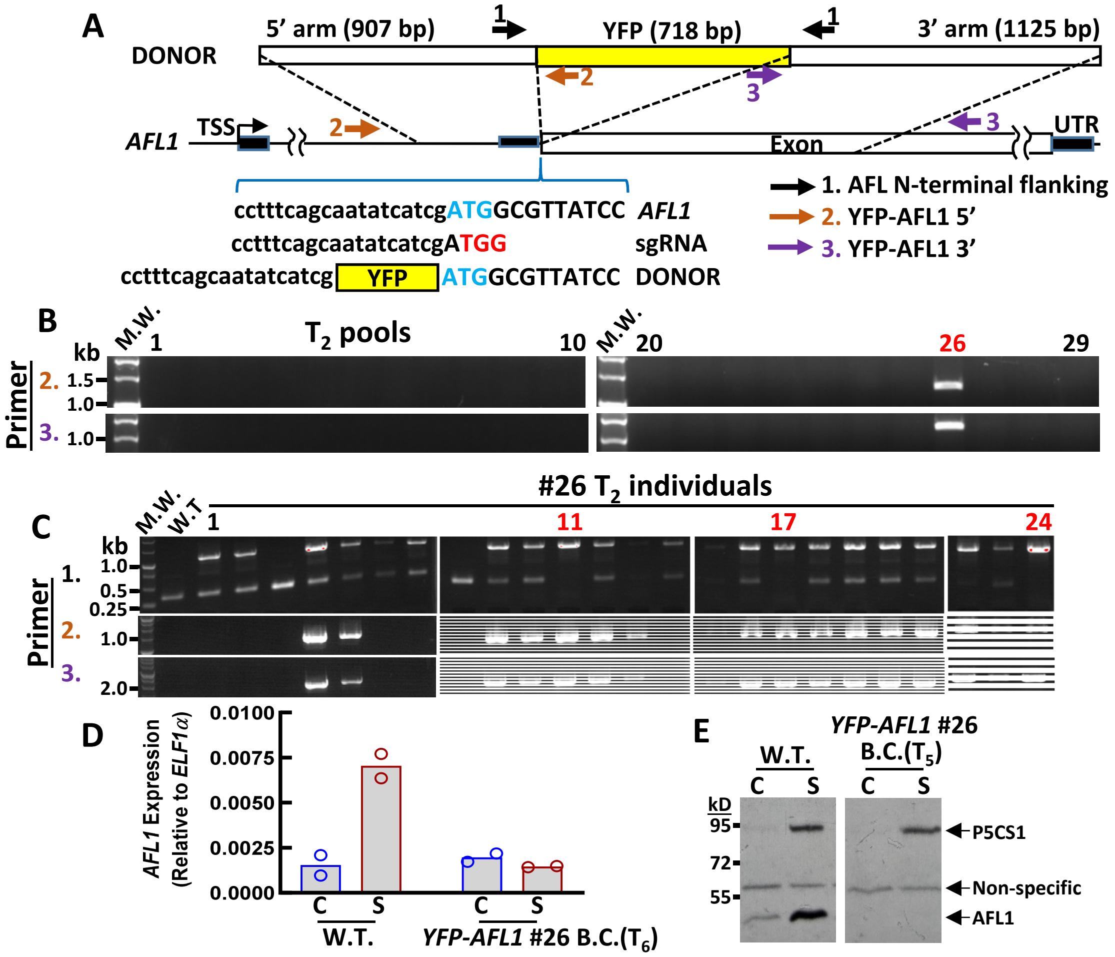


**Figure S14: GT insertion of *YFP* at the 5' end of *AFL1*.**

A. Donor Fragment design and genotyping primer positions for knock-in of N-terminal YFP at the *AFL1* locus. Positions of primers used for genotyping are also shown (primer sequences and amplicon sizes can be found in Supplemental Table I).

B. Genotyping of pooled $T_2$ seedlings from 29 Basta resistant T1 plants using primer sets 2 and 3. Red number indicates T2 seed pool from line #26 with insertion of the YFP donor construct at the *AFL1* locus.

C. Genotyping of individual $T_3$ plants from line 26. Red numbers indicate plants homozygous for the *AFL1-YFP* knock-in.

D. QPCR of *AFL1* in wild type compared to T3 and T6 homozygous plants of *AFL1-YFP* knock-in line #26. Two biological replicates were performed.

E. Immunoblot detection of AFL1. The blot was probed with antisera recognizing both P5CS1 (as a loading control) and AFL1. Protein was extracted from whole seedlings of wild type or T6 homozygous plants collected four days after transfer of seven-day-old seedlings to control or stress (-0.7 MPa) treatments (C = Control, S = Stress). 10 mg of protein was loaded in each lane. All samples are from the same blot with intervening lanes removed for clarity. The predicted molecular weight of AFL1 is 42 kD. The predicted molecular weight of the YFP-AFL1 fusion protein is 66 kD.

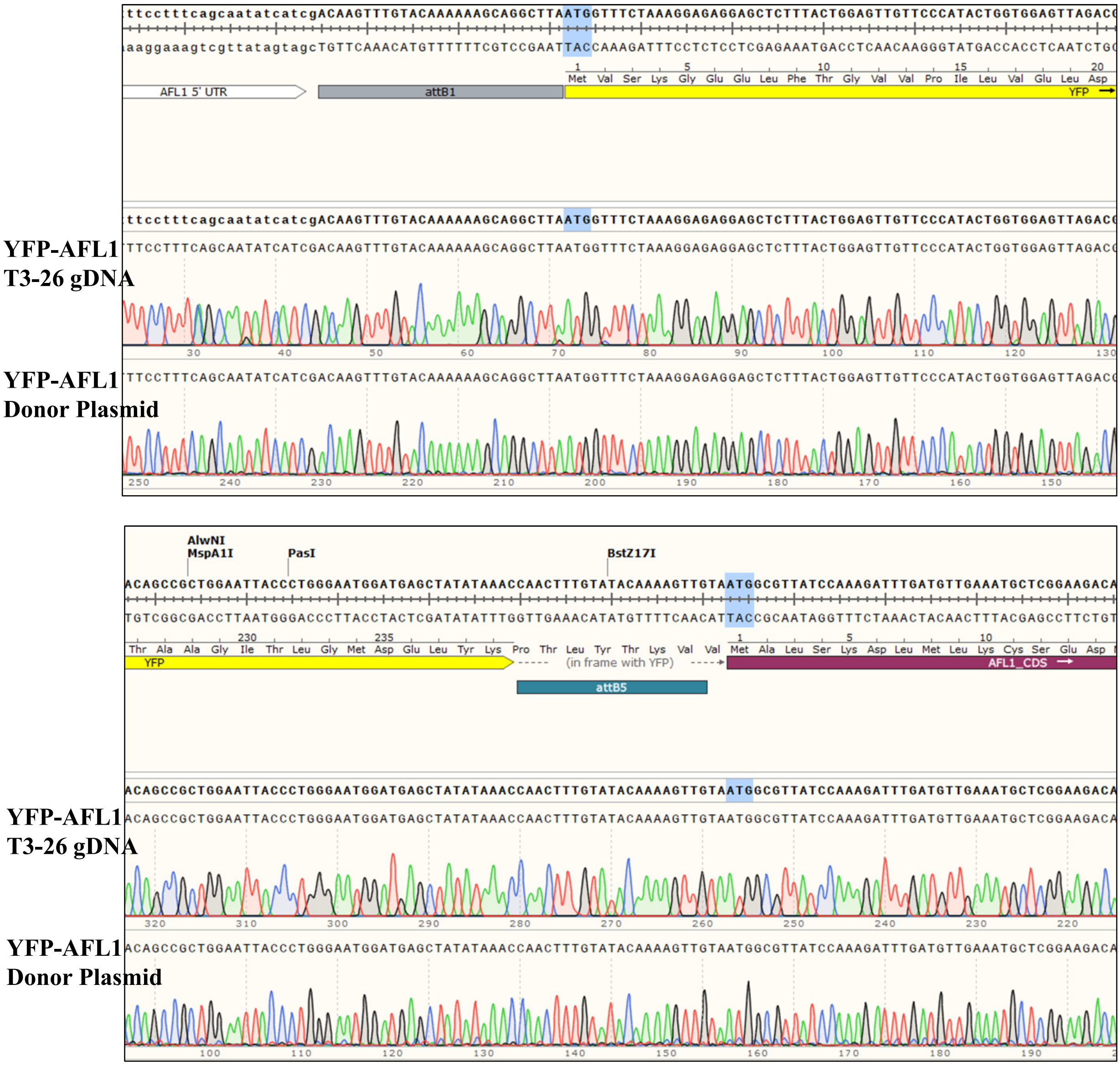


**Figure S15: Sequencing to show the intact and in-frame insertion of YFP at the 5' end of AFL1.**

Sequencing chromatogram of *AFL1* 5' (*YFP-AFL1*) knock-in T3-26 genomic DNA (gDNA) aligned with the sequence of the donor fragment used in the second transformation. Note that this is a different line than the one shown in Figure S2, even though the two lines coincidently had the same number (26) in our labeling of the N-terminal and C-terminal knock-in lines. Alignments were conducted using SnapGene software.

Results show the successful in-frame integration of *YFP*, along with flanking Gateway recombination sequences used to prepare the donor construct, at the 5' end of *AFL1*.
